# DNA tensiometer reveals catch-bond detachment kinetics of kinesin-1, -2 and -3

**DOI:** 10.1101/2024.12.03.626575

**Authors:** Crystal R. Noell, Tzu-Chen Ma, Rui Jiang, Scott A. McKinley, William O. Hancock

## Abstract

Bidirectional cargo transport by kinesin and dynein is essential for cell viability, and defects are linked to neurodegenerative disease. Computational models predict that load-dependent motor detachment strongly determines the outcome of kinesin–dynein tug-of-war, with kinesin-3 and kinesin-2 more load-sensitive than kinesin-1. Yet reconstituted assays show that all three kinesin families compete similarly well against dynein. Previous work demonstrated that vertical forces from optical trapping assays can enhance kinesin-1 dissociation, suggesting that motor behavior may depend strongly on cargo geometry. To measure kinesin detachment and reattachment kinetics under forces applied parallel to the microtubule, we developed a DNA-based tensiometer using an entropic DNA spring linking motors to microtubules. For kinesin-1, -2, and -3, dissociation rates at stall were slower than during unloaded motion, and reattachment kinetics were consistent with a weakly bound slip state preceding detachment. Kinesin-3 behavior further suggested that long KIF1A run lengths arise from multiple short runs connected by diffusive episodes. Stochastic simulations reproduced the measured load-dependent kinetics and enabled direct comparison of transition rates among kinesin families. These results provide insight into how kinesin-1, -2 and -3 transport cargo in complex cellular geometries and compete against dynein during bidirectional transport.

**Impact Statement:** All three kinesin transport families exhibit unexpectedly slow dissociation under load, explaining robust competition with dynein during bidirectional cargo transport.

## Introduction

Bidirectional cargo transport by kinesin and dynein motors is essential for cell viability, and disruptions in transport are linked to neurological diseases including hereditary spastic paraplegia, microcephaly and amyotrophic lateral sclerosis ^1–8^. It has been established that kinesin and dynein, which move in opposite directions along microtubules, are often bound simultaneously to the same cargo ^9–12^. This has led to the ‘tug-of-war’ model, in which the direction of cargo movement is determined by which team of motors dominates ^9,13–17^. How well motors compete is determined by their load-dependent motor properties along with multiple regulation mechanisms, many of which are still emerging ^13,18–21^. Furthermore, the large range of cargo sizes and the complexity of microtubule organization in cells means that motors are subjected to forces both parallel and perpendicular to their microtubule track, which can have differing effects on their mechanochemistry.

Intuitively, a motor’s effectiveness in transporting cargo rests on its ability to remain bound to its microtubule track. Consistent with this, computational simulations have found that the load-dependent off-rate of a motor is the most important determinant of how well a kinesin competes against dynein in bidirectional transport ^22,23^. Single-bead optical tweezers have found that the transport motors kinesin-1, -2, and -3 all act as slip bonds, defined as load accelerating their detachment rate. Their propensity to detach under load varies strongly by family, with relative load sensitivity kinesin-3 > kinesin-2 > kinesin-1 ^24–27^. Based on this behavior, it was surprising that when kinesin-1 was linked to dynein, complexes moved at near-zero speeds for up to tens of seconds, much longer than predicted based on previously measured kinesin-1 off-rates ^25,28,29^. Moreover, kinesin-1, -2, and -3 all fared equally well against dynein, contrary to the differing load-dependent detachment rates measured in single-bead optical tweezer experiments ^30^.

Recent work suggests a solution to this paradox, namely that the ∼micron scale beads used for optical trapping result in significant forces oriented perpendicular to the microtubule as the motor pulls against the force of the trap. First, the load-dependent dissociation rate from single-bead optical trapping was accounted for by a model in which the effects of horizontal loads on detachment is highly asymmetric and vertical loads play a dominant role in detachment, particularly against hindering loads ^31^.

Second, when a three-bead optical trapping geometry was used (in which the motor is raised up on a pedestal bead and the microtubule was held by beads attached to either end of the microtubule) motors remained bound longer than in the single-bead geometry ^24,32^. Third, when kinesin-1 motors were connected to a microtubule by a micron-long segment of DNA, very long residence times were observed, consistent with catch-bond behavior, defined as the off-rate slowing with load ^33^. Fourth, when kinesin-1 was connected to a bead through a micron-long segment of DNA and hydrodynamic forces were imposed on the bead, motor interaction times were insensitive to hindering loads up to 3 pN, indicative of an ideal-bond ^34^. In cells, kinesin and dynein transport cargoes that range from tens of nm in diameter (like vesicles), where motor forces are expected to be aligned parallel to the microtubule, up to several microns (like mitochondria and nuclei), where vertical forces are expected to be much larger. Thus, understanding the influence of vertical and horizontal forces on transport motors is important for understanding the mechanics underlying bidirectional transport in cells.

The goal of the present study was to characterize the load-dependent detachment kinetics of kinesin-1, -2 and -3 motors in a geometry that eliminates vertical forces inherent in traditional optical trapping studies. Building on previous approaches, we used double stranded DNA (dsDNA), which acts as an entropic spring to resist the pN-level forces generated by the motors ^33–38^. We found that kinesin-1, -2, and -3 all remained at stall for multiple seconds before releasing, durations that are substantially longer than the unloaded run times for kinesin-1 and -2. This behavior of slower off-rates under load is defined as a ‘catch bond’, and contrasts with the normal ‘slip bond’ behavior load-accelerated off-rates generally observed for kinesin ^25,39,40^. Following the termination of a stall, motors reengaged with the microtubule with complex kinetics that were consistent with a ‘slip’ state that preceded full detachment. To compare the key transition rates that determined the family-specific motor behaviors, we developed a stochastic model that was able to recapitulate the experimental results for all three motors.

## Results

### Constructing a motor-bound DNA tensiometer

To study motor performance against a resistive load oriented parallel to the microtubule, we constructed a DNA tensiometer consisting of a ∼ 1 μm strand of dsDNA attached to the microtubule on one end and a motor on the other (Figure 1A). We used TIRF microscopy to visualize the motor moving against the entropic elasticity of the DNA spring. Experimental and theoretical studies have established that the force-extension profile of dsDNA is well fit by a Worm-Like Chain model with a persistence length of ∼50 nm ^36,41–43^. As expected from this nonlinear elasticity (Figure 1B), we observed motors moving at near their unloaded velocity to near the ∼ 1 μm contour length of the DNA, at which point they stalled and eventually detached (Figure 1C-E).

**Figure 1.**
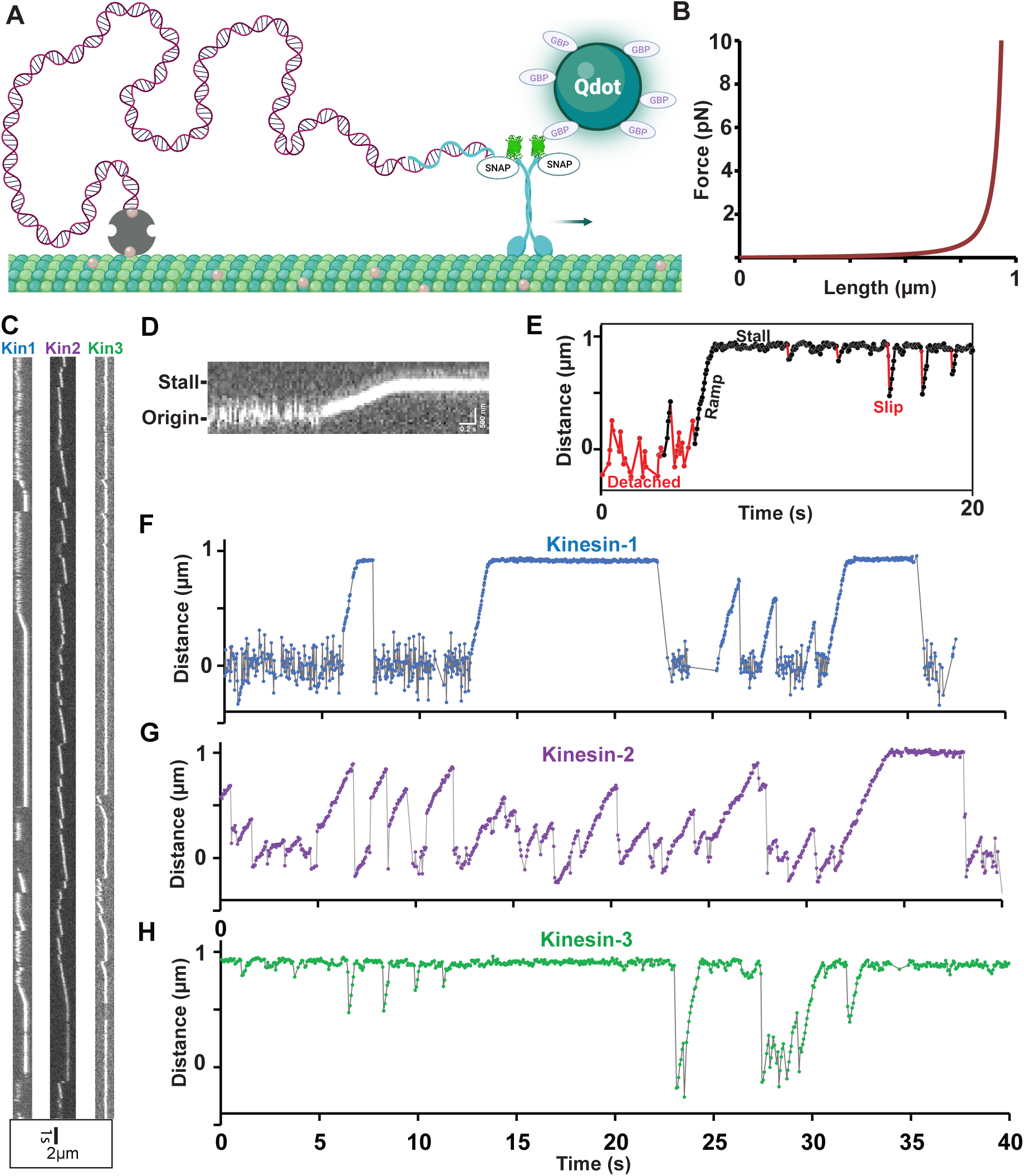
Experimental Design and Raw Data from Motor-DNA Tensiometers. (A) Schematic of a motor-DNA tensiometer, consisting of a dsDNA (burgundy) connected on one end to a kinesin motor through a complimentary oligo (blue), and on the other end to the MT using biotin-avidin (tan and gray, respectively). A Qdot functionalized with GFP binding protein nanobodies is attached to the motor’s GFP tag and used to track motor position. (Not to scale; motor and Qdot are both ∼20 nm and DNA is ∼1 micron) (B) Predicted force extension curve for a worm-like chain 3009 bp dsDNA based on a 50 nm persistence length. We note that our analysis of family-dependent mechanochemistry does not depend on the precise shape of the force-extension curve, only that the motors reach stall. (C) Representative kymographs of motor-DNA tensiometers for kinesins-1, -2 and -3. (D) Enlarged kymograph showing diffusion around the origin, ramp, and stall. (E) Example distance vs. time trace (kinesin-3), highlighting detached durations (red), ramps and stalls (black) where the motor has pulled the DNA taut, and transient slips during stall (red). (F-H) Representative distance vs. time plots for kinesin-1 (F), kinesin-2 (G) and kinesin-3 (H), corresponding to the kymographs in (C). Further examples are shown in Figure 1 – figure supplement 3.

Our DNA-motor tensiometer consists of a 3,009 bp (999 nm contour length) dsDNA ‘spring’ that was synthesized by PCR using a biotinylated forward primer for attachment to the microtubule and a reverse primer containing a 3’ overhang for motor attachment (details in Methods). We investigated members of the three dominant families of kinesin transport motors, kinesin-1 (*Drosophila melanogaster* KHC), kinesin-2 (*Mus musculus* Kif3A), and kinesin-3 (*Rattus norvegicus* Kif1A). The coiled-coil stability and degree of autoinhibition differ between families and can strongly affect motor function ^44–49^. Because our goal was to compare the mechanochemical properties of the motor domains, the head and neck linker of each motor was fused to the stable neck-coil domain (residues 345-406) of kinesin-1, followed by EGFP, a SNAP tag, and His_6_ tag. All of these constructs have been previously characterized by single-molecule TIRF, ATPase, stopped-flow, and optical tweezers studies, as well as by connecting them to a dynein-dynactin-BicD2 complex ^24,30,50–53^. This dimerization approach enables direct comparison to that body of published work and it eliminates the possibility that family-specific differences result from differences in the stability of the neck-coil domain under load or differences in the degree of autoinhibition. Motors were conjugated to an oligonucleotide complimentary to the 3’ overhang of the dsDNA spring via their C-terminal SNAP tag. The DNA tensiometer complex (Figure 1A) was created in a flow cell by sequentially flowing in biotinylated microtubules, neutravidin, biotinylated dsDNA, and Qdot-functionalized motors containing the complimentary oligo (described fully in Methods).

The resulting dsDNA tensiometer kymographs (Figure 1C) show a reproducible behavior of moving, stalling, and returning to origin multiple times, which contrasts with the singular attachment, unidirectional movement and detachment of motors not bound by DNA (Figure 1 – figure supplement 1). Because motors are tethered to the microtubule by the flexible DNA, large fluctuations around the origin are observed when the motor is detached (Figure 1C-H). Consistent with these fluctuations, initial attachment points were variable and roughly normally distributed with a standard deviation of 145 nm (Figure 1 – figure supplement 2). Upon engagement with the microtubule, the motor walks at a steady velocity, consistent with the expected nonlinear stiffness of the dsDNA tether (Figure 1B), until it either disengages or reaches a stall state. Stalls are terminated either by the motor slipping backwards a short distance and restarting a new ramp, or by the motor fully disengaging and returning to the baseline (Figure 1E-H and Figure 1 – figure supplement 3). To confirm that motors are indeed extending the DNA and that Qdots are not enabling multi-motor assemblies, we incorporated Cy5-dCTP into the dsDNA and left the Qdots out of the reaction. In this case, clear extensions of the DNA spring could be observed, and the stall durations were of similar duration (Figure 1 – figure supplement 4). In all subsequent experiments dsDNA was labeled with a low concentration of Cy5-dCTP to confirm colocalization of the DNA and the microtubule before collecting tensiometer data.

### Kinesin-1 and -2 act as catch-bonds at stall

The first question we addressed was: what are the detachment rates of kinesin-1, -2 and -3 motors at stall? The load-dependence of protein-protein interactions can be described as a slip-bond ^54^, defined as a faster off-rate under load; an ideal bond, defined as an off-rate that is independent of load; or a catch-bond, in which the off-rate is slower under load ^39^. Single-bead optical trapping studies consistently find slip-bond characteristics for kinesin-1, 2 and 3 ^25,27,55^, whereas dynein off-rates have been described as a slip-bond or catch-bond ^56–60^.

We define stall duration as the time that a motor stalls against the hindering load of fully extended DNA, without further detectable stepping. Stalls are terminated by the motor detectably (>60 nm) slipping backwards or by disengaging and returning to the origin (Figure 1E). Although we don’t directly measure the stall force, based on the predicted force-extension curve of the dsDNA (Figure 1B), the displacements are consistent with the 4-6 pN stall forces for kinesin-1, -2 and -3 measured using optical traps ^27,61–65^. Stall durations were compared to the unloaded single-motor run durations determined from TIRF kymograph analysis (Figure 1 – figure supplement 1).

To compare unloaded off-rates to off-rates at stall, cumulative distributions of the run and stall durations were plotted for each motor and fit with a single exponential function (Figure 2). The kinesin-1 tensiometer stall duration time constant was 3.01 s, with 95% confidence intervals (CI) of 2.30 to 3.79 s determined via bootstrapping in MEMLET ^66^ with 1000 iterations (N= 78 stalls). In contrast, the kinesin-1 unloaded run duration time constant, measured by a traditional TIRF assay, was 1.04 s, (Figure 2A; Table S1). Stall durations longer than unloaded run durations indicate that load slows the off-rate, the definition of a catch-bond ^39^. Similarly, the kinesin-2 tensiometer stall duration time constant of 2.83 s was longer than its unloaded run duration of 1.07 s, also indicating a catch-bond (Figure 2B). Conversely, the kinesin-3 tensiometer stall duration time constant of 1.89 s was shorter than its unloaded run duration of 2.74 s, indicating a slip-bond characteristic by this definition.

**Figure 2.**
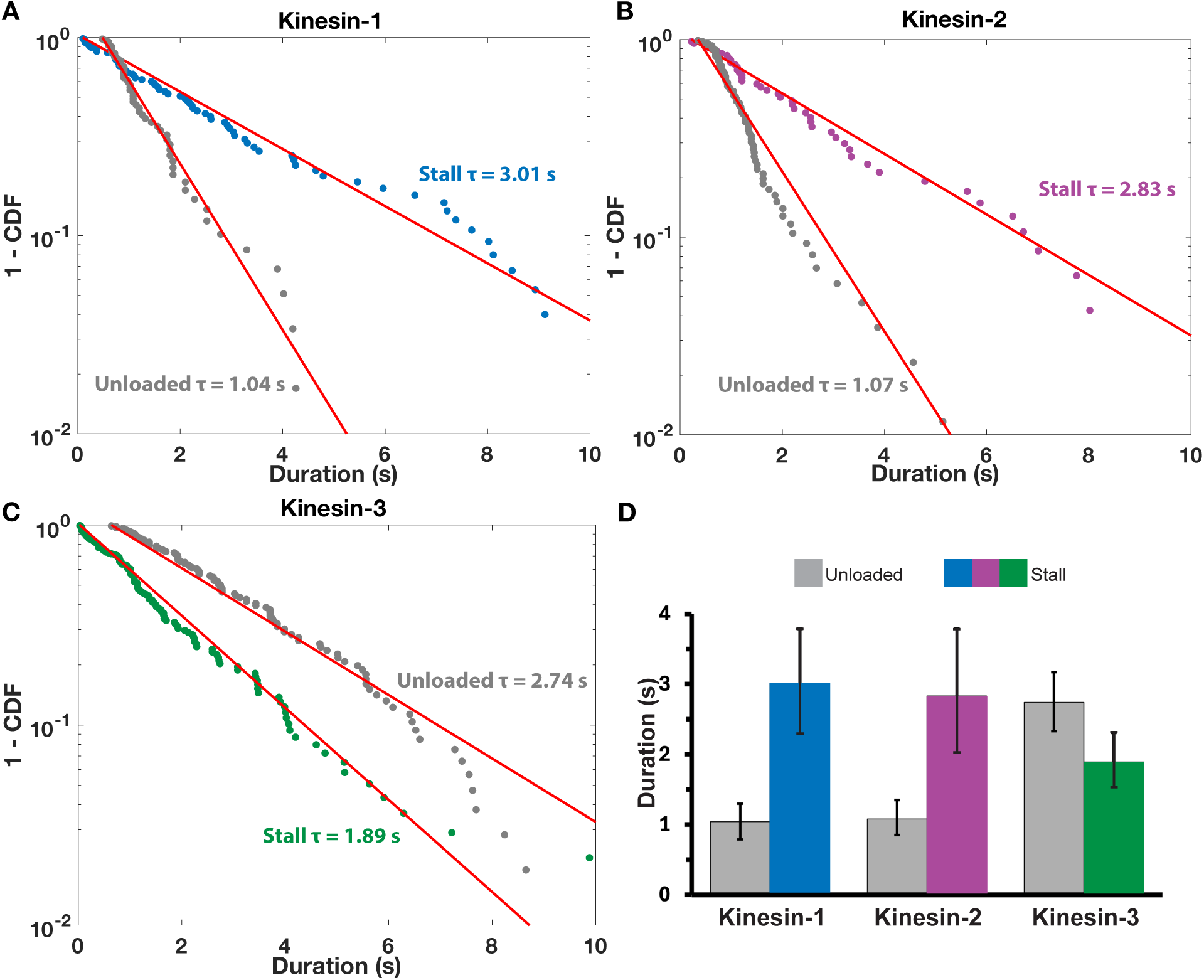
Tensiometer Stall Durations Indicate Catch-bond Behavior for Kinesin-1 and -2. Tensiometer stall durations are plotted for (A) kinesin-1 (blue), (B) kinesin-2 (purple), and (C) kinesin-3 (green). Unloaded run durations for each motor are plotted in gray. Distributions were fit with a single exponential function using MEMLET to generate time constants, representing the mean durations. (D) Comparison of unloaded and stall durations for the three motors, with error bars indicating 95% CI. Stall durations >20s were excluded from the fit (three events for kinesin-1 and two events for kinesin-2). All fit parameters are given in Table S1 and bi-exponential fits of all data including >20 s are shown in Figure 2 – figure supplement 1.

We carried out two additional control experiments. First, to confirm that the neutravidin used to link the DNA to the microtubule wasn’t affecting kinesin motility, we analyzed the run durations of kinesin-1 motors on neutravidin-coated microtubules and found no significant change in the unloaded run duration (Figure 2 – figure supplement 2). Second, we measured the run duration of kinesin-1 linked to a DNA tether that was not bound to the microtubule and thus was being transported (Figure 2 – figure supplement 2). The kinesin-DNA run duration was 1.40 s, longer than the 1.04 s of motors alone (Figure 2A). We interpret this longer duration to reflect the slower diffusion constant of the dsDNA relative to the motor alone, which enables motors to transiently detach and rebind before the DNA cargo has diffused away, thus extending the run duration ^67^. Notably, this slower diffusion constant should not play a role in the DNA tensiometer geometry because if the motor transiently detaches, it will be pulled backward by the elastic forces of the DNA and detected as a slip or detachment event.

### Kinesin-3 detaches readily under low load

To determine whether sub-stall hindering loads affect motor detachment rates, we compared tensiometer ramp durations to the tensiometer stall and unloaded run durations (Figure 3). We defined ramp durations as the time the motor spends walking against the DNA spring before a slip or detachment, or before reaching stall. Although the dsDNA force-extension curve (Figure 1B) predicts negligible loads until the DNA is close to fully extended, there are still non-zero loads imposed during the ramp phase that may affect motor detachment. Based on 10-20% slower ramp velocities relative to unloaded velocities for each motor, we estimated the apparent force to be ∼1 pN (Table S2). To estimate the true detachment rate during the ramp phase in a way that takes into account both the observed detachments and ramps that successfully reach stall, we used a Markov process model, coupled with Bayesian inference methods (detailed in Supplementary Material) to estimate a duration parameter, *τ*, equivalent to the inverse of the detachment rate constant during a ramp. Each increment of time is considered to be an independent opportunity to detach while assuming a constant detachment rate; hence the probability of staying attached to the microtubule through a segment of duration Δ is *e*^-Δ/τ^. Using this method, ramp duration parameters, *τ*, were calculated for each motor, along with 95% credible regions. Finally, to allow for proper comparison, we performed a similar analysis to obtain the stall and unloaded duration parameters along with their 95% credible regions (Figure 3). As expected, the stall and unloaded durations were similar to estimates from curve fitting in Figure 2 (Table S1).

**Figure 3.**
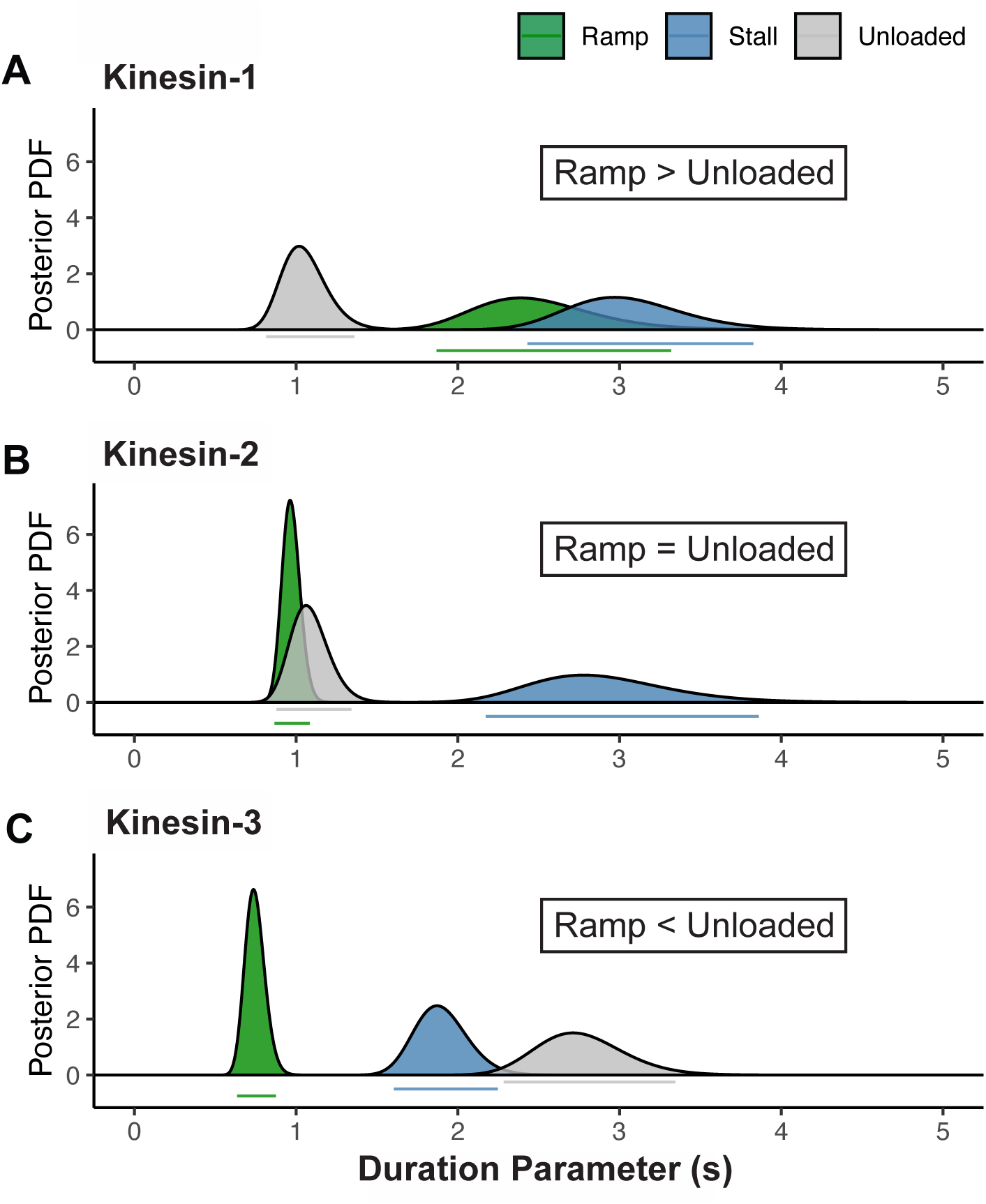
During Ramps, Kinesin-3 Detaches More Readily Than Under Zero Load. Unloaded, ramp, and stall duration parameters were estimated using a Markov process model, coupled with Bayesian inference methods. Curves show the posterior probability distributions of the duration parameters for (A) kinesin-1, (B) kinesin-2 and (C) kinesin-3. Bars below each peak indicate the 95% credible regions for the ramp (green), unloaded (gray) and stall (blue) duration parameters. Notably, the estimated ramp durations are larger, the same, and smaller than the unloaded run durations for kinesin-1, -2, and -3, respectively. For the unloaded and stall durations, this estimation method produces almost identical values as the maximum likelihood estimates in Figure 2 (values provided in Table S1).

The prediction for a slip bond is that against the low loads experienced during ramps, the detachment rate should be equal to or faster than the unloaded detachment rate. This was the case for kinesin-2, where the ramp duration of 0.97 s was within 95% CI of the unloaded run duration of 1.08 s (Figure 3B, Table S1). In contrast, the kinesin-1 ramp duration of 2.49 s was much closer to the stall duration (3.05 s) than the unloaded run duration (1.05 s) (Figure 3A). One possible explanation for the longer kinesin-1 ramp is that the catch-bond character of kinesin-1 engages at low loads rather than rising proportionally to load or engaging only near stall.

The most notable ramp behavior was seen in kinesin-3, where the ramp duration of 0.75 s was nearly four-fold shorter than the unloaded run duration (2.76 s) and was more than two-fold shorter than the stall duration (1.90 s) (Figure 3C, Table S1). As expanded on in the Discussion, the positively charged ‘K-loop’ in the kinesin-3 motor KIF1A is known to interact electrostatically with the negatively charged C-terminal tail of tubulin ^51,68,69^; thus, it is reasonable that even the low loads imposed during ramps are sufficient to overcome these weak electrostatic interactions. The ramp duration is arguably the best definition of the time before KIF1A motors enter a partially dissociated ‘slip’ state, meaning that the observed unloaded durations represent a concatenation of multiple shorter runs interspersed by short diffusive events.

It was notable that the kinesin-3 stall durations at high load are longer than the ramp durations at low load, because this indicates that the kinesin-3 off-rate slows with increasing load. However, because kinesin-3 had the most slip events at stall, we were concerned that there may be undetected slip events below the 60 nm threshold of detection that led to an overestimation of the kinesin-3 stall duration. To test this hypothesis, we plotted the distribution of kinesin-3 slip distances at stall, fit an exponential, and calculated the fraction of missed slip events (Figure 4 – figure supplement 1). From this analysis, we calculated a correction factor of 1.42 that brought the kinesin-3 stall duration down 1.33 s. Notably, this stall duration value is still well above the kinesin-3 ramp duration value of 0.75 s in Figure 3C and thus does not qualitatively change our conclusions.

### Motor reengagement kinetics vary between families

Stall plateaus were terminated by three types of events: 1) small slips that initiated a new ramp, typically within a single frame (∼40 ms), 2) the motor returning to the baseline and reengaging rapidly within a few frames (∼100 msec), or 3) the motor returning to the baseline for a few seconds before reengaging (Figure 4A). We defined a slip event as a displacement of >60 nm from the plateau (distinguishable from normal small fluctuations at stall; Figure 1E) that recovers before reaching within 400 nm of the baseline (outside the range of normal baseline fluctuations; Figure 1F and S2). These slip events have been observed previously for all three motor families in optical trapping experiments and are proposed to represent an intermediate state in which the motor exits the normal stepping cycle but remains associated with the microtubule ^24,70–73^. As an initial analysis, we quantified the fraction of events for each motor (Figure 4A) and found that kinesin-3 had the highest proportion of slip events while kinesin-2 had the lowest proportion. In the context of pulling a large cargo through the viscous cytoplasm or competing against dynein in a tug-of-war, these slip events enable the motor to continue generating force after a small rearward displacement, rather than fully detaching and ‘resetting’ to zero load. Thus, we reanalyzed the stall durations for the three motors where slips are not counted and only disengagements where the motor returns to the baseline are counted as stall termination events (Figure 4B & C, Figure 4 – figure supplement 2). By this definition, stall durations were between 1.5 and 3-fold longer for each motor, including kinesin-3.

**Figure 4.**
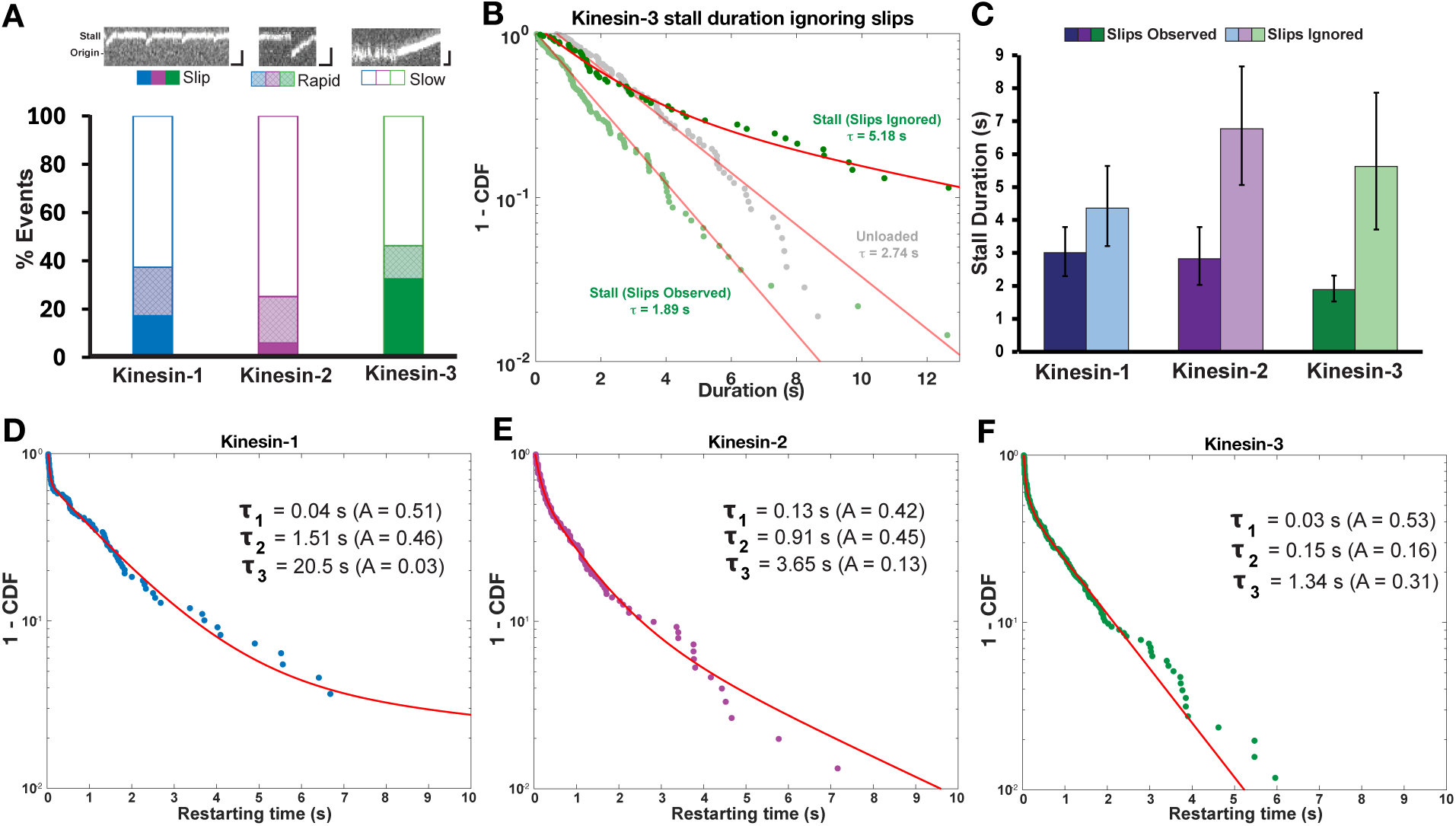
Restart Kinetics for Kinesin-1, -2 and -3. (A) Fraction of slip, fast rebinding, and slow rebinding events for each motor, with example kymographs for each (top; scale bars are 0.5 μm and 0.2 s). Solid colors indicate slips during stall, where the motor resumes a new ramp within a single frame (∼40 ms), crosshatching indicates rapid reattachment events (100 ms) following fall to baseline, and open bars indicate slow reattachment events with >100 ms fluctuations around baseline. (B) Kinesin-3 stall durations, with unloaded run times in gray, stall durations terminated by slips in light green, and stall durations terminated by falling to the baseline (ignoring slips) in dark green. Unloaded and stall durations (replotted from Figure 2) were fit with single exponential functions in MEMLET. Stall durations ignoring slips were fit with a bi-exponential by least squares (ρ1 = 2.01 s [95% CI: 1.52, 2.35 s], A1 = 0.66 s [0.54, 0.83 s], ρ2 = 11.0 s [9.18, 13.70], A2 = 0.33 [0.23, 0.48]). Weighted average of the two time constants is displayed in plot for comparison to other time constants. Similar results for kinesin-1 and -2 are shown in Figure 4 – figure supplement 2. (C) Comparison of stall durations for kinesins -1, -2 and -3 with slips observed as stall terminations or ignored. (D-F) Distribution of restart times for each motor fit to a tri-exponential (least squares). Confidence intervals of parameters determined by bootstrapping with 1000 iterations are given in Table S3.

To obtain a more complete picture of the motor reengagement kinetics for each motor, we plotted the time before starting a new ramp (t_restarting_), including all slips and reattachments (Figure 4D-F). In each case, the distributions included a fast phase and a long tail of slower events. The distributions were fit with a tri-exponential function with the fast phase (30, 130 and 40 msec, respectively) accounting for roughly half of the events (Figure 4 D-F, Table S3). The fast population corresponds to slip and fast reattachment events classified in Figure 4A. The slower phases, which represent detachment events where the motor fluctuated around the baseline before initiating a new ramp, were the fastest for kinesin-3 (time constants of 0.15 and 1.34 s) and the slowest for kinesin-1 (time constants of 1.51 and 20.9 s). Interestingly, the order of the reattachment kinetics (kinesin-3 > kinesin-2 > kinesin-1) and the ∼10-fold ratio of kinesin-3 to kinesin-1 match published bimolecular on-rate constants for microtubule binding from stopped-flow experiments (1.1, 4.6, and 17 μM-1 s-1 for kinesin-1,-2 and - 3, respectively ^50,51,74^).

### Simulating Potential Catch-Bond Mechanisms

To compare the motor detachment and reattachment kinetics between the three kinesin transport families, we carried out stochastic simulations of load-dependent motor stepping, unbinding and rebinding. For simplicity, we reduced the chemomechanical cycle down to a single strongly-bound state (ATP and nucleotide-free states) and a single weakly-bound state (ADP and ADP-Pi states) (Figure 5A). Based on the published load and ATP dependencies of substeps in the kinesin-1 chemomechanical cycle ^70^, we incorporated a load-dependent strong-to-weak transition, k_s-w_. Based on our restarting durations from Figure 4 and previous work ^70–73^, we included both a slip state, from which the motor recovers rapidly, and a detached state associated with a slower recovery. Runs or stalls are terminated by transition from the weakly-bound state into the slip state (k_slip_). Based on backstepping rates observed in optical tweezer experiments, we incorporated a load-independent backward stepping rate of 3 s^-1^, meaning that stall is defined as the load at which forward stepping slows to 3 s^-1^ ^75,76^. For simplicity, we set k_s-w_ = k_w-s_ at zero load, meaning that the motor spends half of its cycle in each state, ^77^ with the rates set to match the unloaded velocity for each motor.

**Figure 5.**
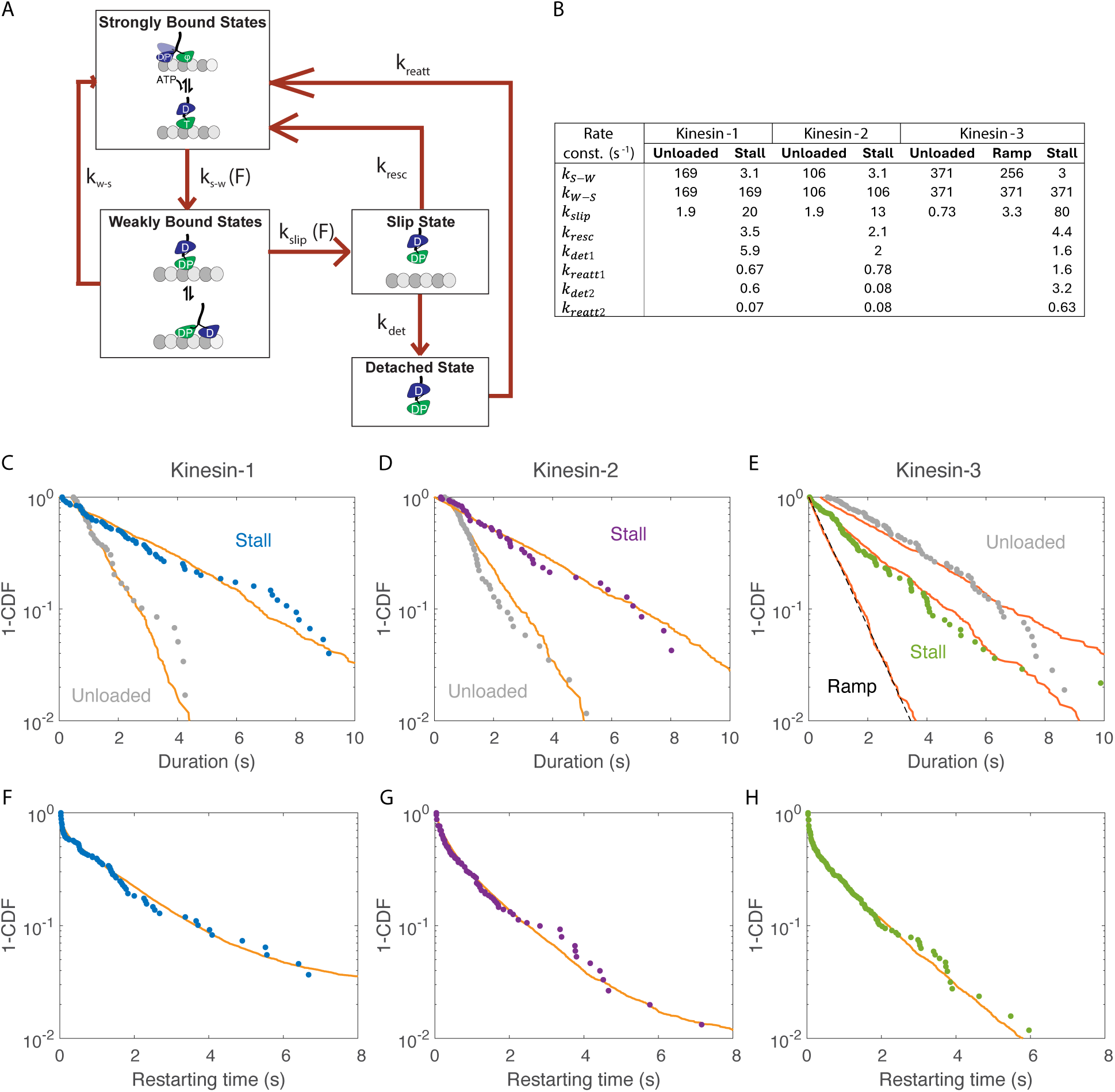
Chemomechanical Model of Proposed Catch-bond Mechanism. (A) Diagram of kinesin chemomechanical cycle model consisting of strongly- and weakly-bound states that make up the stepping cycle, and slip and detached states that terminate runs and stalls. Note that two pathways of detachment from the slip state (and reattachment) are incorporated into the model, but only one pathway is shown for simplicity (see Supplementary Methods for details). (B) Table of rate constants used to simulate unloaded and stall durations and restarting times. All rate constants are derived from fits to experimental data, as described in Supplemental Methods. kS-W and kslip depended exponentially on load 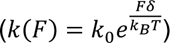 with δ for kS-W of -2.7, -2.4, and -3.6 nm and δ for kslip of 1.6, 1.3 and 2.7 nm for kinesin-1, -2 and -3, respectively; see also Figure 5 – figure supplement 1A). (C-E) Experimental (symbols) and simulated (lines) unloaded and stall durations. 10,000 events were simulated for each condition and plotted with minimum cutoffs matching experiments. Kinesin-3 ramp durations were taken from parameter estimated in Figure 3. (F-H) Experimental (symbols) and simulated (lines) restart times.

Using parameters chosen to match motor behavior under no load (Figure 5B), we were able to reproduce the unloaded run durations for all three motors, as expected (Figure 5C-E). Next, to match the experimental stall durations we incorporated a negative load dependence into the strong-to-weak transition rate (so that stepping slows with load) and a positive load dependence into the transition from the weakly-bound state into the slip state (so slipping occurs more frequently at higher loads). With these parameters (Figure 5B), we were able to reproduce the stall duration distribution for all three motors (Figure 5C-E). Importantly, in this model formulation, dissociation from the weakly-bound state acts as a slip-bond and the kinesin catch-bond characteristics are achieved by the motor spending a larger fraction of its cycle in the strongly-bound state under increasing loads.

To complete our model, we simulated recovery from the slip and detached states. The rate of rescue from the slip state (k_resc_) was set based on the duration and relative amplitude of the fast phase of the restarting times in Figure 4. We posited that detachment follows the slip state, consistent with previous formulations ^70,73^. The two slower restarting time constants from Figure 4 were used to set the reattachment rates, k_reatt_ (see Supplementary Methods for details). Using this approach, we were able to reproduce the restarting durations for all three motors (Figure 5F-H). We note that recapitulating the triexponential restart time distribution in Figure 4D-F required this slip/detached formulation and that lumping all events into a single detached state resulted in single-exponential distribution of recovery times.

The rate constants derived from this model allow for a comparison of the specific transitions that differ between the three motor families. Three features are notable. First, for kinesin-3 the transition into the slip state at stall, k_slip_, is the fastest of the three families, consistent with the observation by eye of the shorter plateaus for kinesin-3 (Figure 1H and 4A) and consistent with slips observed in a previous three-bead optical trapping study ^24^. Second, the rates of rescue from the slip state, k_resc_, for the three motors match within a factor of two. Thus, the fast bimolecular on-rates from stopped flow and the relatively short durations before restarting we observe for kinesin-3 (Figure 4) do not result from a faster reengagement out of the slip state in this model. Third, the slow reattachment rate for kinesin-3 is 10-fold faster than for kinesin-1 and -2. Hence, in this model formulation, both the fast bimolecular on-rates in solution ^51^ and short restart durations observed here for kinesin-3 result from recovery from a detached state rather than rescue from a slip state.

## Discussion

Understanding how motors respond vectorially to external loads is crucial for understanding cargo transport in complex intracellular geometries and how kinesin motors compete against dynein in bidirectional transport ^13,19^. Optical tweezer experiments have provided many essential details of the kinesin mechanochemical cycle under load; however, the bead diameters needed to achieve substantial trapping forces impose vertical forces on the motors. Using DNA as a nanospring enables mechanical experiments using a standard TIRF microscope and allows for simultaneous monitoring of numerous motor-DNA complexes in a single imaging field. With this geometry, a kinesin motor pulls against the elastic force of a stretched DNA nearly parallel to the microtubule, matching the geometry of vesicles measuring a few tens of nanometers. Similar approaches have been used to study myosin, dynein and kinesin-1 in both single-molecule and gliding assays ^33–35,37,38,78,79^. The most striking observation was that members of all three kinesin transport families show catch-bond behavior in which off-rates at stall are slower than those at low or zero loads. Additionally, following disengagement from the microtubule, the three motor families reengaged with the microtubule with complex and family-specific kinetics.

### Comparison to previous work

Despite the clear slip-bond behavior of kinesin-1 seen in single-bead optical traps, there has been growing evidence that kinesin-1 detachment is sensitive to the direction of load. Motor engagement times were shown to decrease when larger beads were employed in single-bead traps, and to be extended in geometries that minimize vertical forces such as the three-bead geometry or when a DNA tether was used ^32,80^. In a study that employed DNA-tethered kinesin-1 to extract tubulin the microtubule lattice, pulling durations of ∼30 s were observed at the lowest motor concentrations, indicative of kinesin-1 catch-bond behavior ^33^. When kinesin-1 was connected to micron-scale beads through a DNA linker and hydrodynamic forces parallel to the microtubule imposed, dissociation rates were relatively insensitive to loads up to ∼3 pN, inconsistent with slip-bond characteristics and instead characteristic of an ideal-bond ^34^. The 3 s kinesin-1 stall duration in our tensiometer falls between the 30 s value for tubulin pulling and the 1.3 – 1.5 s engagement times measured in optical trap and hydrodynamic assays where vertical forces are minimized ^32–34,80^. In contrast to kinesin-1, kinesin-3 (KIF1A) median engagement durations were found to be similarly short in both the one-bead (69 ms) and three bead (62 ms) optical trap geometries ^24^, much shorter than our 1.9 s stall duration in the DNA tensiometer. One difference may be that the millisecond temporal resolution in the optical tweezer enabled detection of many small slips that were undetected by our fluorescence approach; however, extrapolation of our measured kinesin-3 slip distances below our 60 nm detection limit (Figure 4 – figure supplement 1) argues against this. Stiffness differences are an unlikely explanation because at stall the stiffness of the DNA tether (∼4 fold stiffer than optical tweezer) is still sufficiently low to allow for dynamic motor stepping at stall, and in any case it is still below the estimated motor stiffness (see Geometry Calculations in Supplementary methods). One potential explanation is that the microtubule is held under tension in the three-bead experiment, which may alter the lattice properties and affect motor interactions. Finally, it can’t be ruled out that KIF1A is particularly sensitive to vertical loads and small lateral or vertical forces present in the three-bead geometry are absent in the DNA tensiometer geometry. To date there have been no studies on kinesin-2 where vertical loads are minimized, and in single-bead optical traps the off-rate depends strongly on load ^52^.

### Transport kinesins have a catch-bond behavior under hindering loads

What is the mechanism of the observed catch-bond behavior? Cell adhesion proteins such as integrins, selectins, and FimH have been shown to form longer lasting bonds under load, with the proposed mechanisms generally involving an allosteric effect that strengthens the protein: protein interface ^81–83^. However, motor proteins are different in that they cycle in a nucleotide-dependent way between strongly- and weakly bound states, offering multiple potential mechanisms for slower dissociation rates under load. For instance, under a few piconewtons of load, Myosin I was shown to dissociate nearly two orders of magnitude slower than in the absence of load, an effect attributed to load-dependent trapping of ADP in the active site that maintained the motor a high-affinity binding state ^84^. Dynein was also shown to have catch-bond behavior over certain ranges of resisting loads, though the precise mechanism is unclear ^56,57,60,85,86^.

We interpreted our stepping, detachment, and reattachment results using a model that incorporates a load-dependent strong-to-weak transition and a load-dependent entry into a transient ‘slip’ state preceding detachment. The key feature of the model is that under load the motor spends an increasing fraction of its hydrolysis cycle in a strongly-bound state that resists dissociation.

### The role of vertical forces in motor detachment

We next asked whether by considering the different geometries we could reconcile our catch-bond observations with previous single-bead optical tweezer kinesin-1 slip-bond measurements that found the kinesin-1 off-rate increased from 1.11 s^-1^ at zero load to 2.67 s^-1^ at 6 pN ^25^. Using a 440 nm bead diameter and estimated motor length of 35 nm that results in the force being imposed on the motor at a 60° angle (Figure 5 – figure supplement 1B), a 6 pN stall force parallel to the microtubule corresponds to a 10 pN force perpendicular to the microtubule ^25,31^. A model developed by Khataee and Howard was able to fully account for these geometry-dependent off-rates using a two-step detachment process having catch-bond behavior for parallel loads and slip-bond behavior for vertical loads ^31^. Applying that model to our geometry and assuming a purely horizontal load and a 6 pN stall force, the predicted stall duration for kinesin-1 is 77 s, much longer than the 3 s we measure (Figure 5 – figure supplement 1). We approached the detachment process differently, by incorporating a load-dependent transition in the hydrolysis cycle (k_S-W_) and a load-dependent exit from the hydrolysis cycle (k_slip_). To explore effects of vertical forces using our two-state model, we incorporated both horizontal and vertical loads as accelerating detachment from the weakly-bound state, as follows:

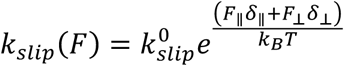

Here F_||_ and F_⊥_ are the magnitude of the parallel and perpendicular loads and δ_||_ and δ_⊥_ represent the distance parameters in each direction (Figure 5 – figure supplement 1). Using δ_||_ = 1.61 nm (Figure 5B), we found that by setting δ_⊥_ = 1.58 nm, we were able to reproduce the slip bond behavior observed in the single-bead optical trap experiments (Figure 5 – figure supplement 1). Notably, this model implies that vertical and horizontal forces have similar effects on the transition rate into the slip state. We stress that this model is a hypothesis that needs further testing. Nonetheless, this is a simple formulation that shows that a motor can display either catch bond or slip bond behavior depending on the geometry of the imposed loads.

### Ramps reveal detachment behaviors at low loads

In addition to reporting on the detachment properties at stall, our DNA tensiometer provides new insights into fast rebinding that occurs during unloaded runs of kinesin-3. It has long been appreciated that the kinesin-3 motor KIF1A achieves long run lengths due to electrostatic attraction between its positively charged Loop-12 (K-loop) and the negatively charged C-terminal tail of tubulin ^45,51,69,87–91^. Furthermore, the KIF1A off-rate in ADP, in which the motor diffuses on the microtubule lattice, was found to match the off-rate during processive stepping in ATP ^51^. The relatively high microtubule affinity of this weakly-bound state suggests that the motor may be undergoing diffusive episodes between processive runs, while maintaining association with the microtubule.

Our DNA tensiometer offers a way to test the hypothesis that the long, unloaded run lengths of KIF1A are due to a concatenation of shorter runs. Due to the nonlinearity of the dsDNA force-extension curve in our DNA tensiometer, the motor is walking against forces below 1 pN for roughly 90% of the distance to stall (Figure 1B). Consistent with this, motor velocities before stall were nearly constant (Figure 1D and S3), and averaged ∼15% slower than unloaded velocities (Table S2), which corresponds to ∼1 pN of force if the force-velocity relationship is linear. Using a Bayesian Inference approach that takes into account motors that dissociate during the ramps as well as those that complete ramps by achieving stall, we measured a nearly four-fold faster KIF1A detachment rate during ramps than under zero load (Figure 3). If, under zero load, the long runs observed were actually a concatenation of a series of shorter runs connected by diffusive weakly-bound events, the diffusive state would likely be unable to withstand even the sub-pN forces from the DNA spring ^90^. For instance, based on a 0.044 μm^2^/s diffusion coefficient (equivalent to a ∼0.1 pN-s/μm drag coefficient ^90,92^), if the motor were in a weakly-bound state for 10 ms, a 1 pN force would pull the motor back 100 nm. Thus, in considering whether KIF1A acts as a catch bond, we used this ramp duration of 0.75 s (Table S1) as the best approximation for the true run unloaded length in the absence of diffusive events.

The ramp durations of kinesin-1 and kinesin-2 also provide insights into how load alters their interactions with microtubules. For kinesin-2, the predicted ramp duration was not statistically different from the unloaded run duration, suggesting that unloaded runs do not include short diffusive episodes. Interestingly the predicted ramp duration for kinesin-1 was nearly the same as the stall duration and much longer than the unloaded duration. One possibility is that the catch-bond effect of hindering load comes into play at low loads and not only at stall where the motor has slowed considerably.

### Motor slips and detachments reflect different processes

Because the DNA tensiometer tethers the motor near the microtubule, such that repeated binding and unbinding events occur, it enables comparison of family-dependent differences in kinesin rebinding kinetics. In addition to clear detachment events, rapid slip and recovery events were observed for all three motors, with highest frequency for kinesin-3 and lowest frequency for kinesin-1. Backwards slipping while maintaining association with the microtubule was first seen for kinesin-8 motors, which are highly processive yet generate only small forces ^71^. Similar backward slips at stall were observed for kinesin-1, with kinetics that suggested a transition such as phosphate release precedes dissociation^73^. Subsequent higher resolution work, enabled by small Germanium nanoparticles, revealed a staircase pattern during these slips with ∼8 nm steps of mean duration 73 μsec, suggesting that the motor was transiently interacting with each tubulin subunit as it slipped backward. Similar slips have also been observed for kinesin-2 and two kinesin-3 family members, KIF1A and KIF1C ^24,72^.

There is some dispute in the literature regarding the kinetics of kinesin-1 recovery from the slip state. Using a single-bead trap, Sudhakar found that 80% of restart events were slips with a time constant of 128 ms ^70^, whereas Toleikis measured slip recoveries that were essentially at the limit of detection (∼1 ms) ^73^. Using a three-bead trap, Pyrpassopoulis measured a 10 ms slip time constant for kinesin-1 ^32^. Our 40 ms slip time constant, which accounts for half of recovery events, is limited by the camera frame rate, and thus is likely an overestimate. In the three-bead geometry, kinesin-3 (KIF1A) slips recovered with a time constant of 1 ms ^24^, faster than the 30 msec (upper limit estimate) we observe. Thus, the precise recovery rate is dependent on the detailed measurement and analysis used. In our DNA tensiometer results, the higher frequency of slips for kinesin-3 relative to kinesin-1 is seen by direct counting (Figure 4A), by the large enhanced stall duration when slips are not counted as termination events (Figure 4C), and by the fast k_slip_ parameter under load in the kinesin-3 model (Figure 5B).

### Catch-bond behavior provides insights into tug-of-war with dynein

In previous simulations of kinesin-dynein bidirectional transport, we found that the strongest determinants of kinesin’s ability to compete against dynein were the load-dependent motor dissociation rate and the motor rebinding rate ^22,23^. Simply put, if motors detach, then the opposing motor wins. The finding here that all three dominant kinesin transport families display catch-bond like behavior at stall necessitates a reevaluation of how motors function during a tug-of-war. There is evidence that dynein forms a catch bond or at least an ideal (load independent) bond ^14,56,57,60,85,86,93^; thus, kinesins and dyneins are primed to strongly oppose one another.

The catch bond results here help to explain previous in vitro work in which one kinesin and one dynein were connected through a complementary ssDNA ^14,28,30^. It was found that the motor pairs had periods of near-zero velocity that lasted for many seconds, considerably longer than kinesin’s unloaded off-rate. Furthermore, kinesin-2 and kinesin-3 also showed these sustained slow tug-of-war periods despite their reported faster load-dependent off-rates from optical tweezer studies. The functional catch-bond behavior observed here provides a simple explanation for these sustained kinesin-dynein stalemates.

Importantly, the load-dependent off-rates of both kinesins and dynein are expected to depend on the cargo geometry. A 30 nm vesicle would lead to forces on the motor nearly parallel to the microtubule surface, whereas when transporting a micron-scale mitochondria the vertical forces would be larger than the horizontal forces. Cargo geometry and stiffness are also expected to play a role; for instance, deformation of a cargo, either due to compliance of the cargo or to multiple motors pulling on it will tend to reduce vertical force components on the motors. Additionally, any ‘rolling’ of a spherical cargo following motor detachment will tend to suppress the motor reattachment rate. The present work emphasizes that along with motor type, motor number, motor autoinhibition, and the growing list of regulatory proteins, the geometry with kinesin and dynein engage in a tug-of-war can be an important determinant of the speed and direction of cargo transport in cells.

## Methods

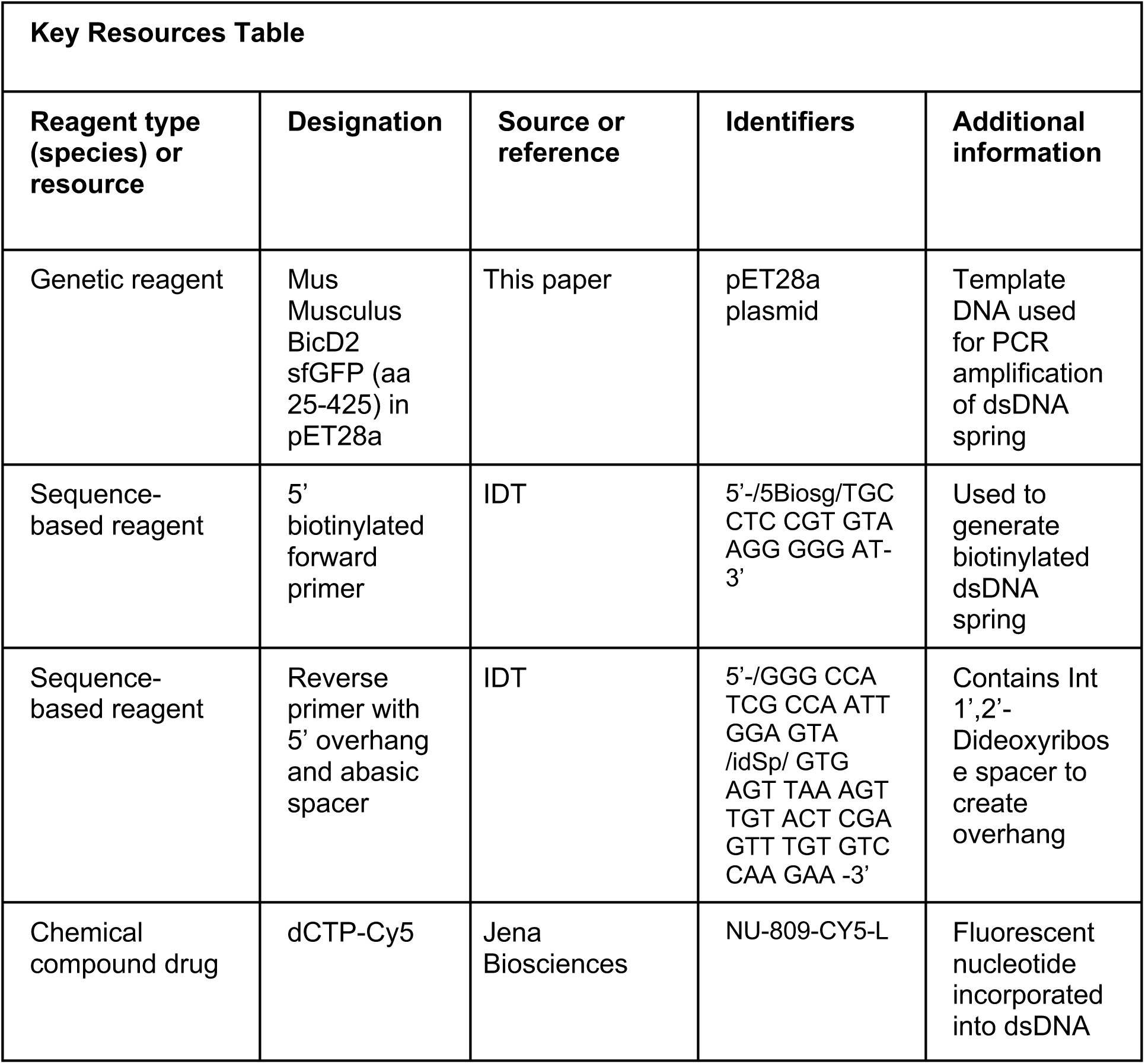
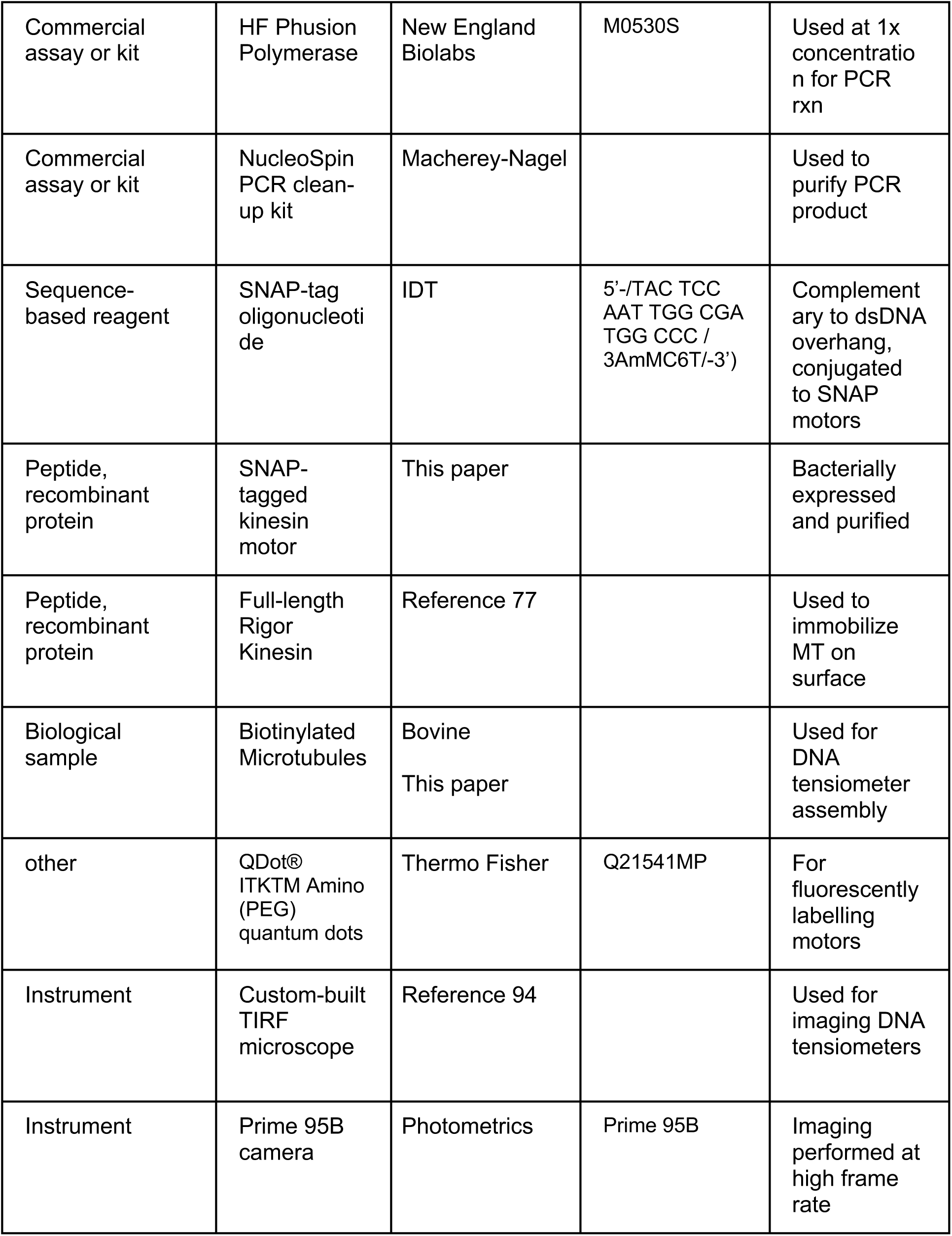
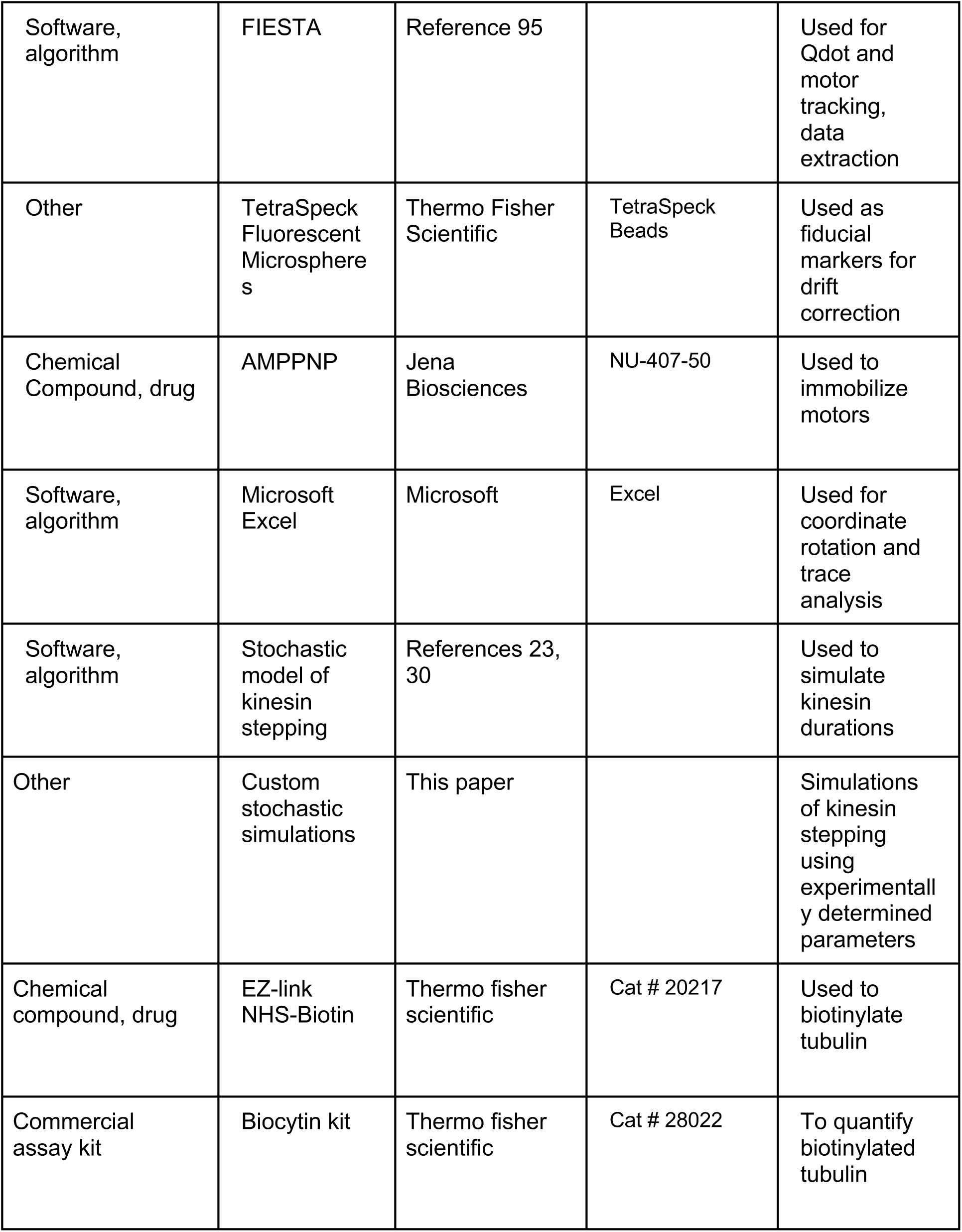
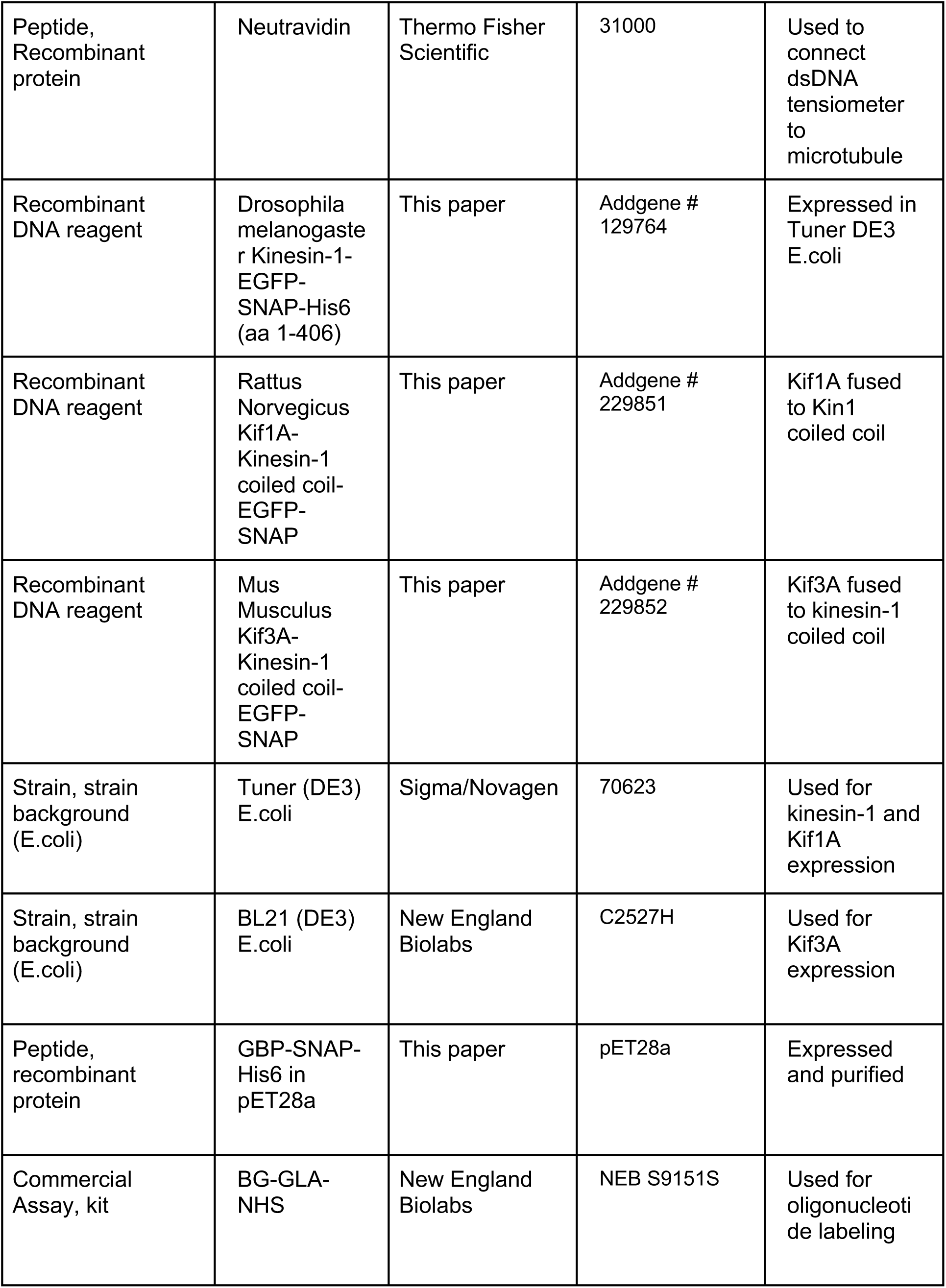

### DNA Tensiometer Construction

For the dsDNA spring, a 5’ biotinylated forward primer (5’-/5Biosg/TGC CTC CGT GTA AGG GGG AT-3’) and a reverse primer with a 5’ overhang (5’-/GGG CCA TCG CCA ATT GGA GTA /idSp/ GTG AGT TAA AGT TGT ACT CGA GTT TGT GTC CAA GAA-3’) were used to create a 3009 bp dsDNA by PCR from plasmid *Mus Musculus* BicD2-sf-GFP (aa 25-425) in pet28a. The abasic Int 1’,2’-Dideoxyribose spacer (idSp) creates an overhang by terminating the polymerase. All oligonucleotides were purchased from IDT. Each 50 μL PCR reaction contained: 1x Phusion HF buffer, 198 μM dNTPs, 2 μM dCTP-Cy5, 0.5 μM primers, 3 ng template DNA and 1 U/50 μL HF Phusion Polymerase. Fluorescent dsDNA used in Figure 1 – figure supplement 4 had 10 μM dCTP-Cy5 and 190 μM dNTPs. The PCR reaction was carried out in a thermal cycler with the following procedure: 98℃ for 30 s, then 45 cycles of 98℃ for 10 s, 58℃ for 30 s and 72℃ for 1.5 min, then lastly 72℃ for 5 min. The product was purified using a NucleoSpin® PCR clean-up kit and the concentration determined by absorbance on a Nanodrop 2000c Spectrophotometer. DNA bands were visualized on a 1% agarose gel with ethidium bromide staining.

### Motor-Microtubule-Tensiometer Assembly

Motors were bacterially expressed, purified and linked through its SNAP tag to an oligonucleotide (5’-/TAC TCC AAT TGG CGA TGG CCC / 3AmMC6T/-3’) complementary to the dsDNA overhang. Details of motor expression, purification and labeling, as well as tubulin biotinylation and polymerization are given in Supplementary Information. The DNA tensiometer was assembled on the microtubule as follows. The following three buffers are made on the same day of the experiment: C2AT (BRB80, 10 μM Taxol, 2 mM MgATP, 2 mg/mL Casein), 2AT (BRB80, 10 μM Taxol, 2 mM MgATP) and Imaging Solution (BRB80, 10 μM Taxol, 2 mg/mL Casein, 2 mM MgATP, 20 mM D-glucose, 0.02 mg/mL Glucose oxidase, 0.008 mg/mL Catalase, 0.5% BME and 2 mg/mL BSA). Full-length rigor kinesin was used to attach microtubules to the coverglass ^77^.

Tensiometers were created in the flow cell using the following work flow: C2AT, 5 min > Rigor kinesin, 5 min> C2AT wash > BioMT, 5 min > 2AT > 8 nM Neutravidin, 5 min> 2AT > 10 nM Bio-dsDNA-Overhang, 5 min> C2AT > 4 nM KinesinMotor + 40nM Qdot-GBP (pre incubated in tube on ice for >15 min) in imaging solution, 10 min > Imaging solution wash. Note that because casein can contain free biotin, casein-free 2AT buffer was used during avidin-biotin binding steps. Following assembly, the Qdot connected to the motor was imaged on a custom-built TIRF microscope, described previously ^94^. Raw data were typically collected at 25 fps (range of 20-40 fps) on a Photometrics Prime 95B camera.

### Data Analysis

Movies were uploaded into FIESTA software ^95^ and Qdot intensities were tracked using a symmetric 2-D gaussian function to obtain x,y,t data for each Qdot. When drift correction was needed, TetraSpeck™ Fluorescent Microspheres (Thermo) and immobile Qdots were used as fiducial markers. The smallest position errors at stall in FIESTA fitting were 3-4 nm, which matched the positional error of Qdot-labeled motors stuck to microtubules in AMPPNP. Points with position errors greater than 20 nm were excluded because they often involved clearly spurious position estimates. Notably, many tensiometers had small segments of missing data due to the Qdot fluctuating out of the TIRF field or blinking; these occurred most often during periods when the motors were detached from the microtubule.

After obtaining X and Y positions of linear motor tracks in Fiesta, we rotated and translated the data in Excel to generate X versus t traces. The apparent origin was determined by averaging the points where the motor is fluctuating on its tether. In rare instances where no fluctuation was observed, the approximate origin was calculated by averaging the starting positions of all the ramps within the tensiometer (Figure 1). We then measured the ramp time, distance traveled, stall durations, reattachment times and starting positions. Tensiometers occasionally ended with the Qdot signal going dark, denoting either bleaching or failure of the Qdot-motor or motor-DNA connection. Notably, no clear instances of motor-Qdots walking past the plateau point (denoting the tensiometer breaking) were observed. Stalls that terminated due to the tensiometer going dark or the video ending were excluded from analysis.

### Stochastic Modeling

Kinesin run and stall durations were simulated by using a modified version of published stochastic model of kinesin stepping ^23,30^. A motor is either in a strongly-bound state or a weakly-bound state (Figure 5A). At each timepoint, a motor in the strongly bound state can transition into the weakly-bound state with a first order transition rate constant, *k_S_*_-*W*_ or step backward by 8 nm with a constant rate, *k_back_* = 3 *s*^-1^. A motor in the weakly-bound state can complete an 8-nm forward step by transitioning back to the strongly-bound state with rate constant, *k_W_*_-*S*_, or it can disengage from the microtubule with transition rate, *k_slip_*.

For simplicity, we set *k_S-W_* = *k_W_*_-*S*_ at zero load. Load-dependent transition rates were defined as:

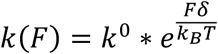

where *k*^0^ is the unloaded transition rate, *δ* is the characteristic distance parameter and *k_B_T* is the Boltzmann’s constant multiplied by the absolute temperature, equal to 4.1 pN-nm at 25° C. Stall force was set to 6 pN. Unloaded and stall durations were simulated by starting the motor in strongly-bound state and continuing until it transitioned into the slip state, with 1000 simulations for each condition. Restart times were simulated by starting the motor in the slip state. From there, the motor can reengage by transitioning into the strongly-bound state or switch to one of two detached states having either a slow or fast recovery rate. Restart simulations were run for 10,000 iterations. Model parameters were constrained by experimental data, described fully in Supplementary Methods.

## Acknowledgments

This work was originally conceived as part of a NIH-funded collaborative modeling project to Will Hancock, Scott McKinley, John Fricks, and Peter Kramer (R01GM122082). We thank Qingzhou Feng, Scott Pflumm, and Adheshwari Ramesh for early efforts on this project and all members of the Hancock Lab for helpful discussions. This work was funded by NIH Grant R35GM139568 to W.O.H.. C.R.N. was supported by NIH postdoctoral fellowship F32GM149114, R.J. was supported by NIH Training Grant T32GM108563, and S.A.M. was supported by the NSF-Simons Southeast Center for Mathematics and Biology (SCMB) through grant NSF-DMS1764406 and Simons Foundation-SFARI 594594.

## Data Availability

Data for all figures and code used in simulations are publicly available at http://doi.org/10.26207/98h3-5e15.

## Supporting Information

## Supplementary Methods

### Biotinylated MT

Tubulin was purified from bovine brain as previously described ^1,2^. Biotinylated tubulin was made by polymerizing microtubules for 45 minutes at 37℃, adding a 12-fold excess of EZ-link NHS-biotin (Thermo Fisher 20217), incubating at 37℃ for another 30 minutes, and then pelleting. Microtubules were then depolymerized, polymerized, and pelleted twice to obtain pure biotinylated tubulin. Tubulin concentration was measured by absorbance, and the fraction biotinylated measured using the Biocytin Kit (Thermo Fisher 28022).

For DNA tensiometer experiments, microtubules were polymerized with 10% biotinylated tubulin and functionalized by adding 8 nM neutravidin to the flow cell, making 10% the upper limit of biotin-neutravidin coated tubulin in the microtubules. However, this percentage is likely lower due to: a) less than 100% labeling in the biotinylation reaction, b) the potential for preferential incorporation of unlabeled tubulin over biotinylated tubulin in the polymerization reaction, and 3) incomplete occupation of biotinylated tubulin by neutravidin. Based on Fig. 3C of Korten and Diez ^3^, the small reduction in the unloaded run duration for kinesin-1 that we measured on biotinylated microtubules (Fig. S6) corresponds to a ∼2% biotinylation ratio. Thus, that is our best estimate for the fraction of biotin-neutravidin functionalized tubulin. A second argument that neutravidin was not acting as a roadblock in the DNA tensiometer experiments was the fact that ramp durations were not consistently shorter than unloaded run durations on control microtubules lacking neutravidin (Fig. 3).

### Motor Expression, Purification and Oligo Conjugation

*Drosophila melanogaster* Kinesin-1-EGFP-SNAP-His_6_ (aa 1-406) was expressed in Tuner (DE3) E.coli (Addgene #129764). Cells were grown in Terrific Broth (Sigma Aldrich) at 37℃ with shaking at 180 rpm for 4-6 hours until an OD of greater than 1 was reached, then induced with 120 mg IPTG and shaken overnight at 21℃. Cells were harvested the next day, pelleted, resuspended with 1x PBS, frozen, and stored at −80℃. *Rattus norvegicus* Kif1A (aa 1-351)-Kif1A neck linker (NL) (17aa)-Kinesin1 coiled-coil (aa 345-406)-EGFP-SNAP was expressed in Tuner (DE3) E.coli similarly to kinesin-1 (Addgene #229851). *Mus musculus* Kif3A (aa 1-342)-Kif3A NL (17aa)-Kinesin1 coiled-coil (aa 345-406)-EGFP-SNAP was expressed in BL21(DE3) E.coli (Addgene # 229852). Kif1A and Kif3A constructs were synthesized into the kinesin-1 construct backbone by GenScript. His_6_ tagged GBP-SNAP in pet28a was also expressed and purified similarly ^4–7^.

Bacterial cell pellets (from 800 mL culture) were thawed and motors were purified via Ni affinity chromatography as described previously ^4,8,9^. Motor proteins were eluted in a buffer containing 20 mM phosphate buffer, 500 mM sodium chloride, 500 mM imidazole, 10 μM ATP and 5 mM DTT. The concentration of pre-labeled motors was then measured by absorbance at 488 nm (using the EGFP extinction coefficient 55,900 M^-1^cm^-1^), and proteins were visualized with SDS PAGE.

Amine-terminated oligonucleotides (IDT) were resuspended and desalted into 200 mM sodium borate buffer, and the concentration measured by absorbance. The desalted oligo was then mixed with 20-fold excess of BG-GLA-NHS (NEB S9151S, dissolved in DMSO) in 100 mM sodium borate and 50% DMSO and incubated at RT for 30 min. The mixture was then desalted into 1x PBS buffer (containing 1 mM DTT and 1 mM MgCl_2_). The elution profile was measured by absorbance and the fractions of BG-oligo were pooled. The pre- and post-labeled oligos were visualized on a 10% TBE-Urea gel and stained with SYBR green I. Excess BG-oligo was stored at -20℃.

Immediately following Ni column purification of motors, BG-oligo was mixed with the eluted motor at a 5:1 ratio and incubated on ice for 1 hr. The mixture was then diluted with 1x PBS + 1 mM MgCl_2_ sufficient to reduce the imidazole concentration to below 80 mM, and a second Ni-affinity purification was carried out to remove the excess BG-oligo. The protein was eluted in the same elution buffer and flash frozen in liquid N_2_ in the presence of 10 μM MgATP, 1 mM DTT and 10% sucrose. Final protein concentration was measured by EGFP absorbance. SDS-PAGE and native PAGE were used to estimate the percentage of motors that have an oligo conjugated to them, typically ∼50% were labeled. Oligo-labeled motors were kept at -80℃ for up to a year.

### Qdot Labeling

For labelling GFP motors, Qdots were functionalized with GFP-binding protein (GBP) as follows. QDot® ITKTM Amino (PEG) quantum dots (Thermo Fisher Q21541MP) were buffer exchanged by transferring 250uL into a 100K ultrafiltration unit and adding 1x PBS pH 7.4 to make up the filter volume of 4 mL. The sample was centrifuged to the original volume of 250 μL before more buffer was added and the process was repeated 3x. The Qdots were then transferred to a glass vial, BG-GLA-NHS was added in 50-fold excess in a 100 mM sodium borate buffer containing 50% DMSO (v/v), and the reaction incubated for 1 hr at room temperature on a rotator. Excess BG-GLA-NHS was removed by carrying out 5 complete buffer exchanges with a 100K centrifugal concentrating filter. The concentration of BG-Qdots was determined on a plate reader based on a calibration curve from the initial Qdot stock. BG-Qdots were then mixed with GBP-SNAP at a 1:50 ratio and incubated on ice for 1 hour. Qdot-GBP was stored at 4℃ for up to 6 months. On the day of an experiment, Qdot-GBP was mixed with GFP labeled motors at a 10:1 ratio to prevent multi-motor Qdots and incubated on ice for at least 15 minutes before visualization by TIRF.

### Geometry Calculations

The maximum theoretical vertical forces in the DNA tensiometer are calculated as follows. The quantum dot does not affect the vertical forces because it is attached to the GFP on the kinesin independent of the attachment to the DNA tether. The kinesin head is ∼5 nm ^10^ and the kinesin neck-coil (residue 345-392 at 0.15 nm /residue) is ∼7 nm. Assuming that the entire neck-coil is sticking up, the DNA tether starts ∼12 nm above the microtubule surface. If the motor and the ∼5 nm neutravidin ^11^ are on the same side of the microtubule, then there is a 7 nm difference relative to the 960 nm extension of the DNA (0.3 degree angle), meaning vertical forces are < 1% of horizontal forces. However, there is evidence that the neck-coil lays parallel to the microtubule ^12^, and so this difference is likely smaller than this. If the motor and the neutravidin are on opposite sides of the microtubule, then the vertical force component (12 nm motor + 5 nm neutravidin + 25 nm Mt; 2.5° angle) will be <5 % of horizontal; however, the direction of load will be pointing into the microtubule surface (and oriented laterally) rather than oriented away from the microtubule. Thus, the geometry is nearly optimal to minimize any upward vertical forces on the motor.

Another consideration when comparing the DNA tensiometer to optical trap measurements is the relative stiffness of the trap and dsDNA. Optical traps stiffnesses are generally in the range of 0.05 pN/nm ^13,14^. To calculate the predicted stiffness of the dsDNA spring, we computed the slope of theoretical force-extension curve in Fig. 1B. The stiffness is highly nonlinear and is <0.001 pN/nM below 650 nm extension. We compare motor performance under this low stiffness regime to the unloaded case in Fig. 3. In contrast, at the predicted stall force of 6 pN (960 nm extension), the dsDNA stiffness is ∼0.2 pN/nm, which is stiffer than most optical traps, but it is similar to the estimated 0.3 pN/nm stiffness of kinesin motors themselves ^13,14^. An 8 nm step at the 0.2 pN/nm stiffness of the dsDNA leads to a 1.6 pN jump in force and at the 0.05 pN/nm stiffness of an optical trap leads to a 0.4 pN jump in force; this is important because it means that in both cases the motors are likely dynamically stepping at stall. Because both experimental approaches allow for dynamic stepping at stall and because the stiffnesses of the instrument in both cases are less than the motor stiffness, there is no reason to expect that differences in stiffness between optical traps and the dsDNA spring lead to different motor detachment kinetics.

### Fitting Equations

Data in Figures 2, S5 and S6 were fit using MEMLET ^15^ to the following single exponential function:

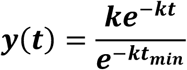

where *k* is the rate constant (inverse of the time constant) and *t_min_* is the minimum cutoff of the distribution.

Data in Figure S4 were fit using MEMLET to the following bi-exponential function:

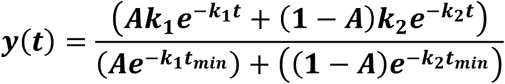

Here A is the amplitude of the first phase, *k_1_* and *k_2_* are the rate constants (inverse time constants) of the two phases, and *t_min_* is the minimum cutoff of the distribution. This approach corrects the amplitudes for missed events, which can differ for the two phases Cumulative distribution data in Figure 4 were fit by least squares in Matlab to the bi-exponential function:

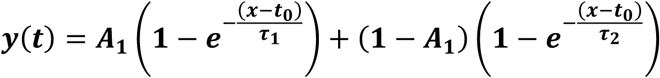

Here ***τ*_1_** and ***τ*_2_** are time constants (inverse rate constants) of the two phases, A_1_ is the amplitude of the first phase, and t_0_ is the minimum cutoff time of the distribution.

Amplitudes are normalized to account for missed short events as follows:

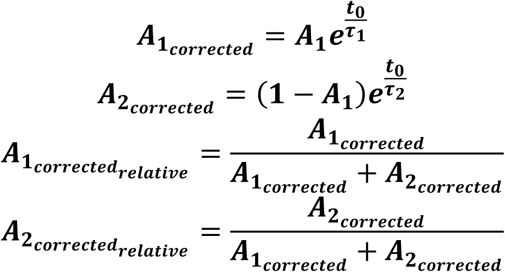

### Inference Strategy for Ramp Duration Parameter

The primary challenge in assessing ramp runs is that these segments can end for one of two reasons: either (1) the motor detaches, or (2) the motor reaches a distance sufficiently far from the anchor to enter a stalled state. By contrast, unloaded runs and stall segments all end in detachments from microtubules. So, while taking a simple average of segment durations will result in an “average time until detachment” for unloaded and stalled segments, this is not the case for ramps. Moreover, ambiguity as to when a ramp run begins further confirms that simply averaging ramp segment durations will lead to errors. This issue is similar to the one raised in Rayens et al. ^16^ in which the authors sought to estimate the average stationary segment length for motor-lysosome complexes in vivo, even though the length of some stationary segments exceeded the length of the observation window.

One way to address the problem of truncated durations is to assume that state-switching satisfies the Markov property, which is to say that whether an agent switches states in a given time step is independent of states and switches that occurred during previous time steps. for continuous time Markov chains, models are expressed in terms of rates, and so the natural quantity of interest here is the detachment rate, which might commonly be denoted *k*_off_. Within the text, we wish to compare ramp state properties to unloaded and stall states which are quantified in terms of their average duration. In this note we therefore write the detachment rate in terms of a duration parameter *τ* = 1/*k*_off_. Generically, the probability that a continuous-time Markov chain does not change state in a segment of time [*a*, *b*] is given by the formula:

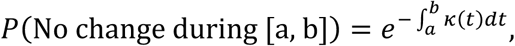

where *κ*(*t*) the (possibly evolving) rate at which changes occur. If we assume that the detachment rate 1/*τ* is constant, then in any segment of length Δ, the probability that a motor does not detach is

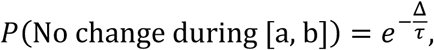

Suppose that (*t*_0_, *t*_1_,…, *t_n_*) are observation times of a tensiometer run and let *T* denote the time that a motor would detach were it not for transitions to stall. Then the probability that a detachment occurs in the interval [*t_k_*, *t_k_*_+1_) is product of the probabilities it does not detach in all preceding segments multiplied by the probability it does detach in the final one. This idea is shown in the diagram below. The initial green segments are ramps which may result in a detachment (yellow star), or result in conversion to a stall state (red line) which always ends in a detachment. In describing our approach to inference, we denote the ramp durations 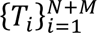 and the stall durations are 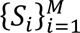. This notation scheme implies that there were *N* detachment events during ramp segments.

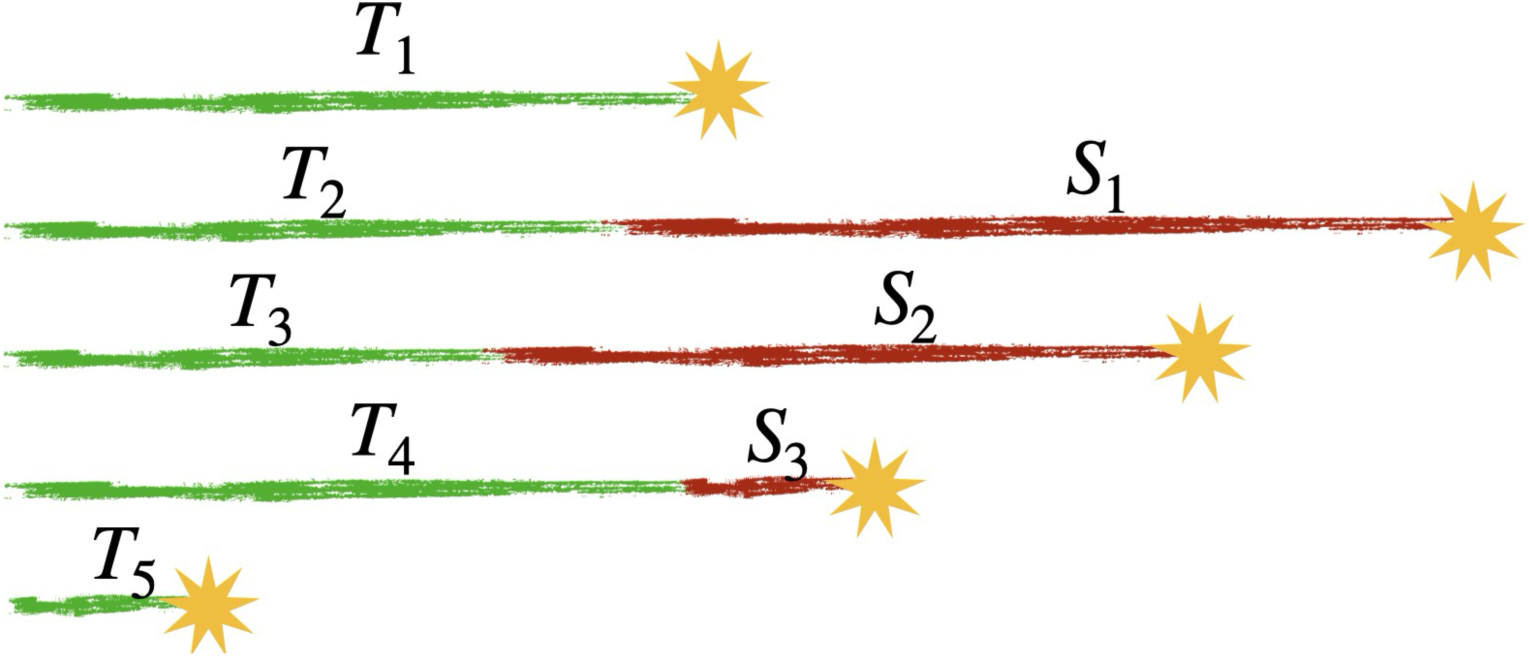

Mathematically, we write

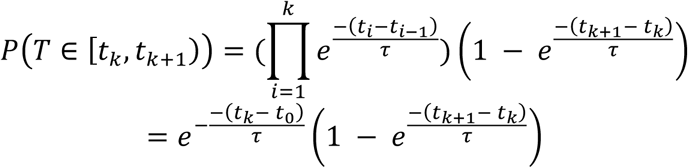

Assuming that time increments are evenly spaced and of size Δ, and also assuming that Δ is small compared to the duration parameter *τ*, we can take the first term of the Taylor expansion of the (*t_k_*_+1_ *− t_k_*) term and simplify:

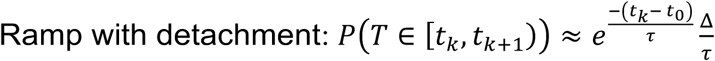

On the other hand, suppose that the motor switches to a stalled state after time *t_k_*. Then the initial product of non-detaching segments remains the same, and final term is removed:

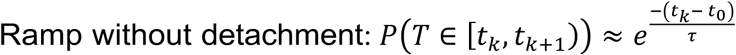

To unify the notation, let 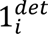 denote the event that the *i* ramp run detached before reaching stall phase. We have

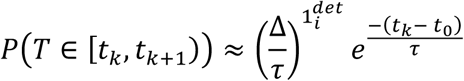

To perform inference on multiple runs, we assume they are independent and so, again, we can take a simple product. As described in the caption of Figure 1, let 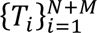 be an enumeration of ramp durations, where *N* is the number of ramps that detached and *M* is the number of ramps that reached a stall state. It follows that the likelihood of observing a set of trajectories *X* given a duration parameter *τ* can be written

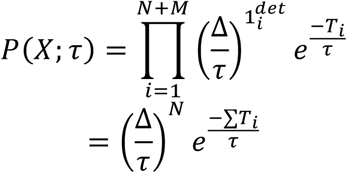

From a Bayesian perspective, we can use this likelihood function to create a posterior distribution for the duration parameter *τ*. In this work we have used a scale-free uninformative prior ^17^ of the form *π*(*r*) = *r*^-1^. Together with the likelihood, we have that the posterior distribution has the form

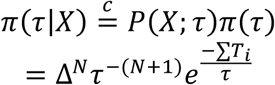

where *c̳* means “equals up to a constant depending only on *X”*. Looking at the factors that depend only on *τ* this is an Inverse Gamma distribution.

In this way, we reach the conclusion that if *T* is the total time spent in the ramp states and if *N* is the number of detachments while in the ramp state, the posterior distribution for the duration parameter is

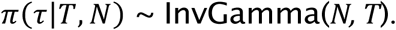

This means that it has pdf

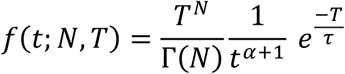

and, most importantly for the purpose of estimation, the mean of the posterior distribution is simply *T*/*N*. In other words, the simple Bayes estimator for a collection of ramp segments is:

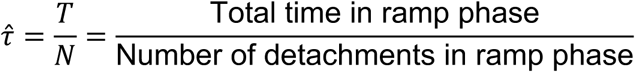

To set our 100(1 *− α*) %-credible regions, we used the middle-*α* probability range from the posterior distribution. This means that the 95% credible region is defined as the interval between the 0.025-quantile and 0.975-quantile of the posterior distribution. These regions are indicated as solid lines segments beneath the posterior distributions depicted in Figure 3 in the main text.

### Compensating for Difficulty in Observing Short Runs

For unloaded and ramp runs there is a significant risk that short runs will be unobserved. For ramp runs, short ramp phases might be masked by the fluctuations that occur when the motor is detached. Meanwhile, short, unloaded runs might not be recognized among many motors in a wide field of view. Among kinesin-1 runs, for example, the shortest recorded unloaded run was 0.48s, despite the frame rate being significantly smaller. The minimum recorded times, *t*_min_ (seconds), for each category of run observations is as follows:

|  | Stall | Ramp | Unloaded |
| --- | --- | --- | --- |
| Kinesin-1 | 0.1 | 0.14 | 0.48 |
| Kinesin-2 | 0.22 | 0.17 | 0.348 |
| Kinesin-3 | 0.03 | 0.03 | 0.641 |

The likelihood function therefore needs to be adjusted to be conditioned on being greater than the minimum observation time for that cohort. Continuing with the notation from above (for convenience taking *t*_0_ = 0, let *T_i_* denote the *i*th run duration. Then the joint likelihood function is therefore

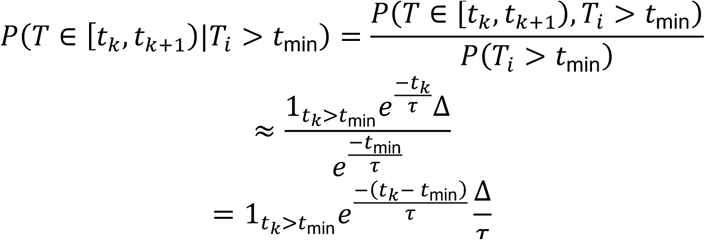

For a collection of paths *χ*, the likelihood function becomes

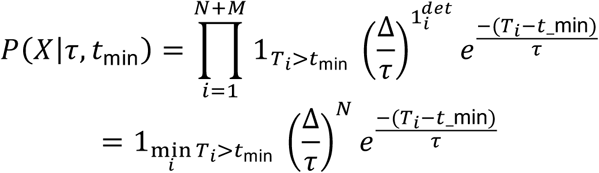

Note that for any fixed value *τ*, the maximum of the likelihood function over all *t*_min_ values is 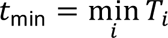. Rather than constructing a joint Bayesian posterior for the pair (*τ*, *t*_min_), we simply adopted the maximum likelihood value for *t*_min_ within each cohort and proceeded as described above with the likelihood function

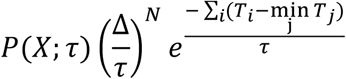

### Rate Constant Constraints in Stochastic Model

To constrain the rate constants in the stochastic model simulations using the experimental results, the model was solved analytically, as follows. The forward stepping rate and the load-dependent velocity are:

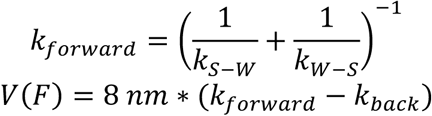

Ignoring the relatively slow backstepping rate of 3 s^-1^ in the stepping cycle, the bound duration is equal to the duration of each step multiplied by the predicted number of steps before detaching:

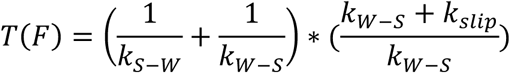

The forward stepping rates for unloaded kinesin-1, -2, -3 simulations were set based on the unloaded velocities to 84.5, 53 and 151.5 steps per second, respectively. At stall, defined as a force of 6 pN, the forward stepping rates were set to 3 s^-1^, matching the backstepping rate. The transition to the slip state, *k_slip_*, was chosen based on the run and stall durations for each motor.

To simulate restarting times, the rate constants were constrained by the restart time constants (*τ*) and respective weights (A) in Figure 4. The rescue rate (*k_resc_*) and two slip-to-detached transition rates (*k_det_*_1_& *k_det_*_2_) were determined based on the shortest time constant (*τ*_1_) as follows,

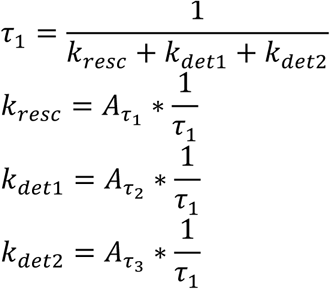

This formulation accounted for the relative amplitudes of the three time constants. The reattachment rates (*k_reatt_*_1_ & *k_reatt_*_2_) were defined based on the two slower restart time constants by considering that the time to restart is the sum of time spent in the slip state plus the time spent in the detached state. The two reattachment rates were defined as follows,

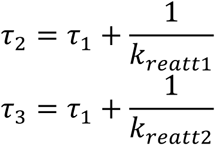

This process was repeated for the three motor families.

---

**Table S1.**
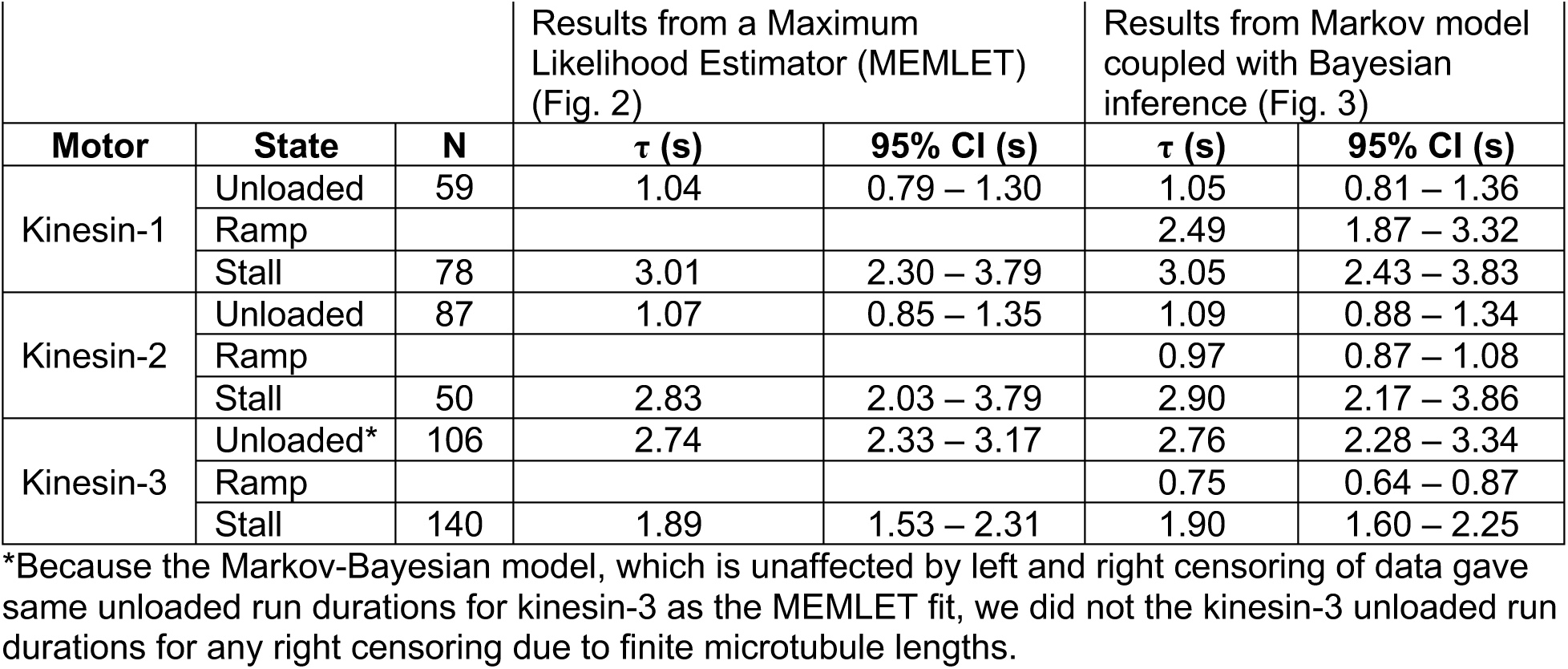
Fit results for unloaded, ramp, and stall durations.

**Table S2.** Predicted force imposed on the motor during the ramp phase.

| | $V_{\text{unloaded}}$ (nm/s) | $V_{\text{ramp}}$ (nm/s) | $V_{\text{ramp}} / V_{\text{unloaded}}$ | $F_{\text{predicted}}$ (pN)* |
| --- | --- | --- | --- | --- |
| Kinesin-1 | 651 | 572 | 0.88 | 0.72 |
| Kinesin-2 | 401 | 324 | 0.81 | 1.14 |
| Kinesin-3 | 1458 | 1187 | 0.81 | 1.14 |

\*Based on linear force-velocity relationship with a 6 pN stall force

**Table S3:** Reattachment Statistics.

| Reattachment Tri-Exponential Statistics |  |  |  |  |  |  |
| --- | --- | --- | --- | --- | --- | --- |
| | $\tau_1$ (s) | $A_1$ | $\tau_2$ (s) | $A_2$ | $\tau_3$ (s) | $A_3$ |
| Kinesin-1 | 0.04<br>[0.03, 0.04] | 0.51<br>[0.34, 0.74] | 1.50<br>[1.35, 1.65] | 0.46<br>[0.38, 0.58] | 20.5<br>[0.00, 61.13] | 0.03<br>[0.00, 0.07] |
| Kinesin-2 | 0.13<br>[0.11, 0.16] | 0.42<br>[0.00, 1.00] | 0.91<br>[0.45, 1.00] | 0.45<br>[0.14, 1.00] | 3.65<br>[0.00, 8.61] | 0.13<br>[0.00, 1.00] |
| Kinesin-3 | 0.03<br>[0.02, 0.03] | 0.53<br>[0.32, 0.84] | 0.15<br>[0.09, 0.21] | 0.16<br>[0.11, 0.25] | 1.35<br>[1.24, 1.44] | 0.31<br>[0.24, 0.41] |

\*95% Confidence intervals determined by bootstrapping with 1000 iterations are indicated in brackets

**Figure 1 – figure supplement 1.**
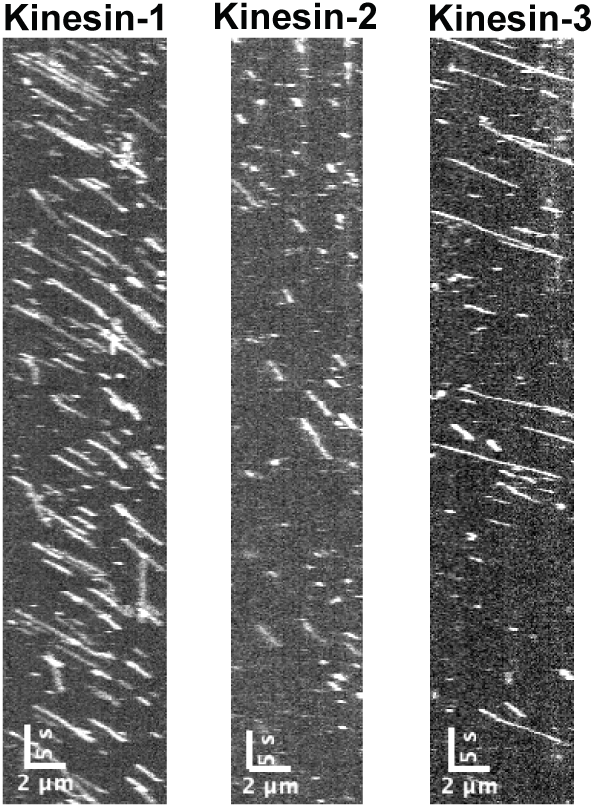
Kymographs of unloaded GFP-labeled kinesins. GPF-labeled motors conjugated to their complimentary oligo were visualized via TIRF at a concentration of 1 nM at 5 fps. No neutravidin, Qdot or DNA are present in these unloaded controls. A fraction of kinesin-3 unloaded run durations were limited by the length of the microtubules, but fitting to a model that took into account missed events gave a similar mean duration as an exponential fit, and so no correction was made (Table S2).

**Figure 1 – figure supplement 2.**
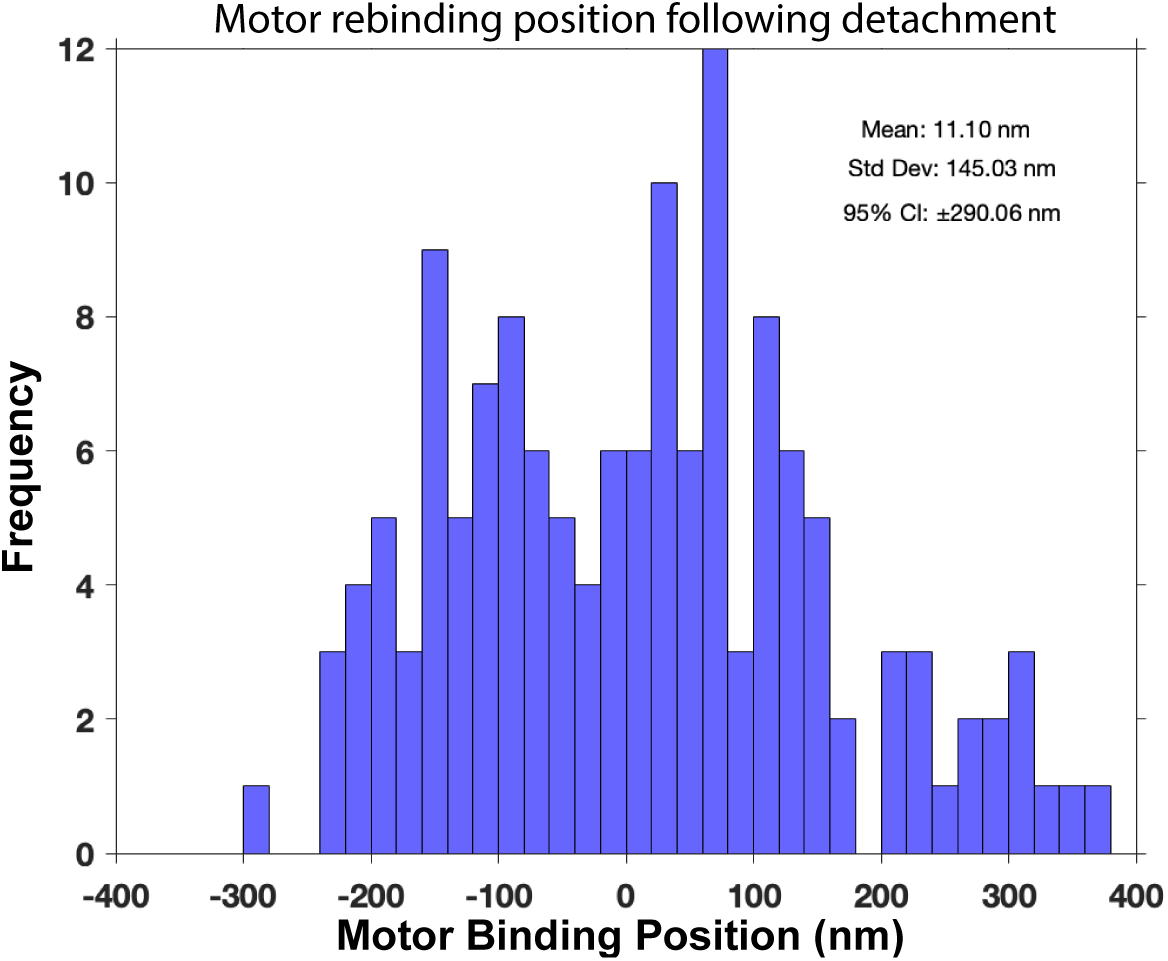
Distribution of initial motor binding positions. The initial positions were measured for events where the motor was clearly dissociated from the microtubule and fluctuated around origin on its DNA tether (N=141). The zero position was determined as the center point around which the detached motor fluctuated. The initial motor binding position was determined by the first start point of a ramp. The width of the gaussian distribution, quantified by the SD, demonstrates the large search space of the motor attached to the flexible ∼1-micron dsDNA tether. The mean of +11 nm likely results from some of the motors moving before the first frame acquisition, giving a small positive bias. The larger population seen at >+200 nm relative to <-200 nm may result from dissociation of motors that bind under assisting loads (negative displacements) and strengthening of motors that bind under hindering loads (positive displacements). Data are from kinesin-1 and kinesin-3 tensiometers.

**Figure 1 – figure supplement 3.**
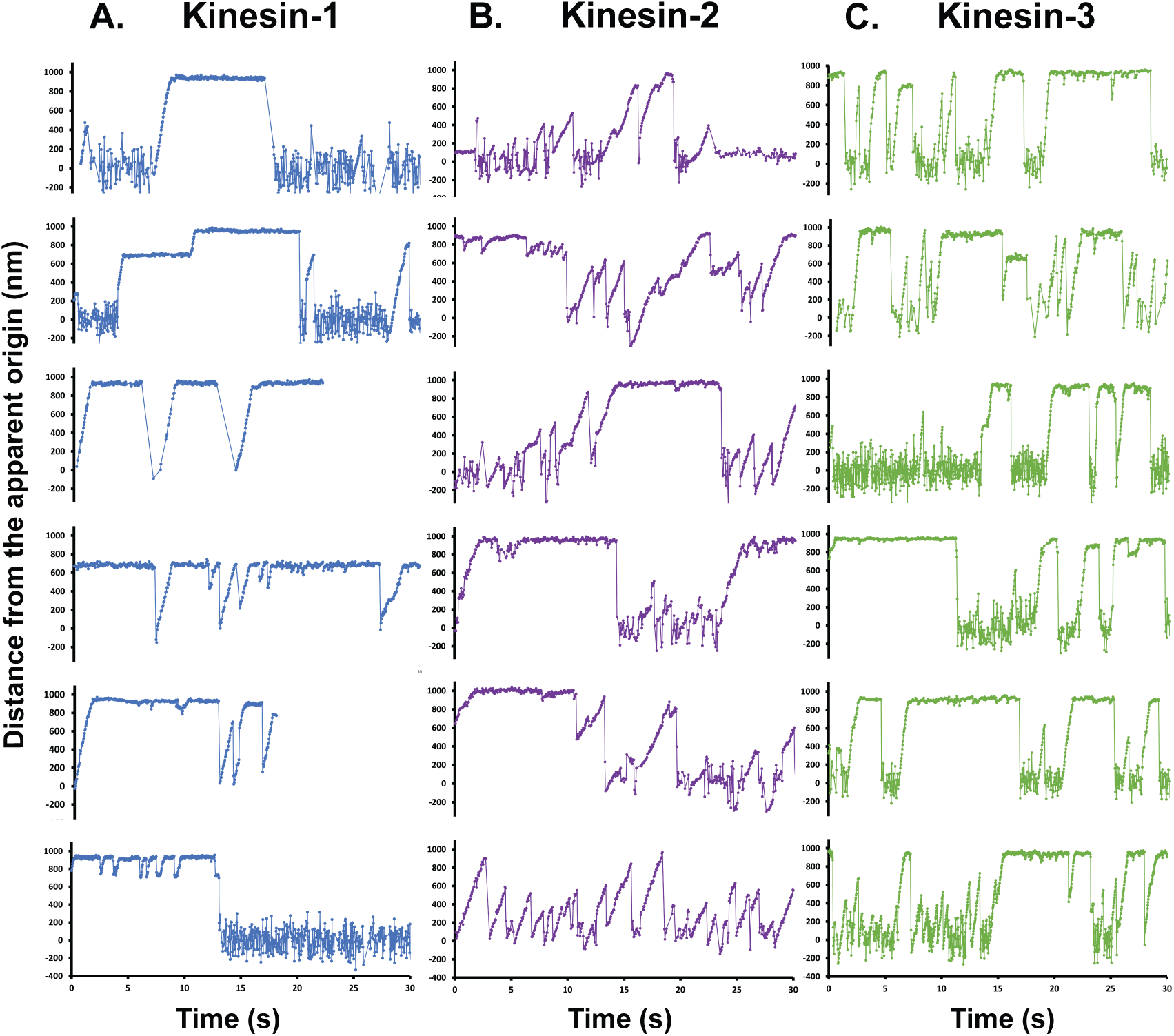
Further kinesin tensiometer examples. Distance versus time plots of (A) kinesin-1, (B) kinesin-2 and (C) kinesin-3 traces. Notably, some stalls are very stable, whereas other (particularly for kinesin-2 and kinesin-3) show fluctuations, presumably due to small slips and backstepping at stall. Other features to note include pauses in the motile segments, small changes in velocity, and repeated ramps for kinesin-2. Roughly 20% of tensiometers extended less than the expected 1 μm distance, stalling repeatedly at 500 nm or 800 nm. These apparently shorter DNA strands may result from DNA secondary structures or from primer binding at a secondary sequence. After in-depth comparison of the data we found that the stall durations, reattachment rates, and starting positions were unaffected by the shorter DNA length and thus included the shorter tensiometers in our data set.

**Figure 1 – figure supplement 4.**
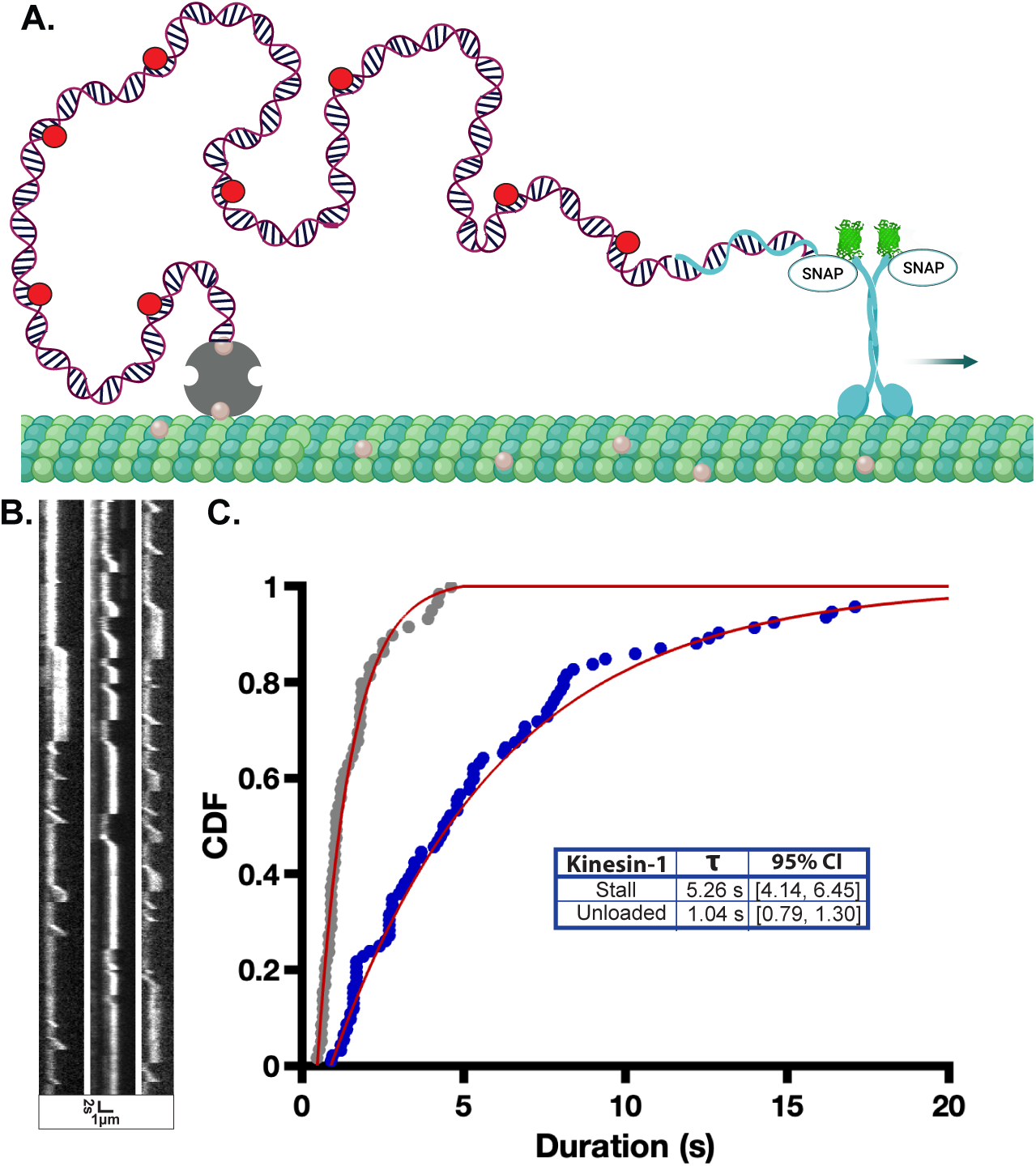
Long stall durations are observed in the absence of Qdots. Because the Qdots used in our experiments are functionalized with multiple GBP nanobodies, there is the possibility that the long stall durations observed were caused by multiple motors bound to the Qdots. To test this, we ran a control experiment where, instead of labeling the motors with Qdots, we fluorescently labeled our dsDNA by incorporating 5% dCTP-Cy5 during the PCR reaction to create fluorescent dsDNA. By removing Qdots from the system, the potential for multimotor events is eliminated. (A) Diagram of control experiment. (B) TIRF kymographs of representative bright kinesin-1 DNA tensiometers collected at 5 fps, showing the typical extension and stall profiles we observed in Figure 1, with the difference that the entire dsDNA is visualized rather than the Qdot. (C) CDF plot of the fluorescent DNA tensiometer stall durations of kinesin-1, fit with a single exponential function using a maximum likelihood estimator (MEMLET). Importantly, kinesin-1 continued to have much longer stall durations than its unloaded run durations (gray points with fit; reproduced from Fig. 2A), ruling out multi-motor interactions as the cause of the long stall durations. The longer stall durations here (5.26 s) compared to the Qdot stall durations (3.01 s; Fig. 2A) is attributed to the 5 fps frame rate used in here, which makes it more difficult to detect short slip events that are observed with the 25 fps Qdot movies.

**Figure 2 – figure supplement 1.**
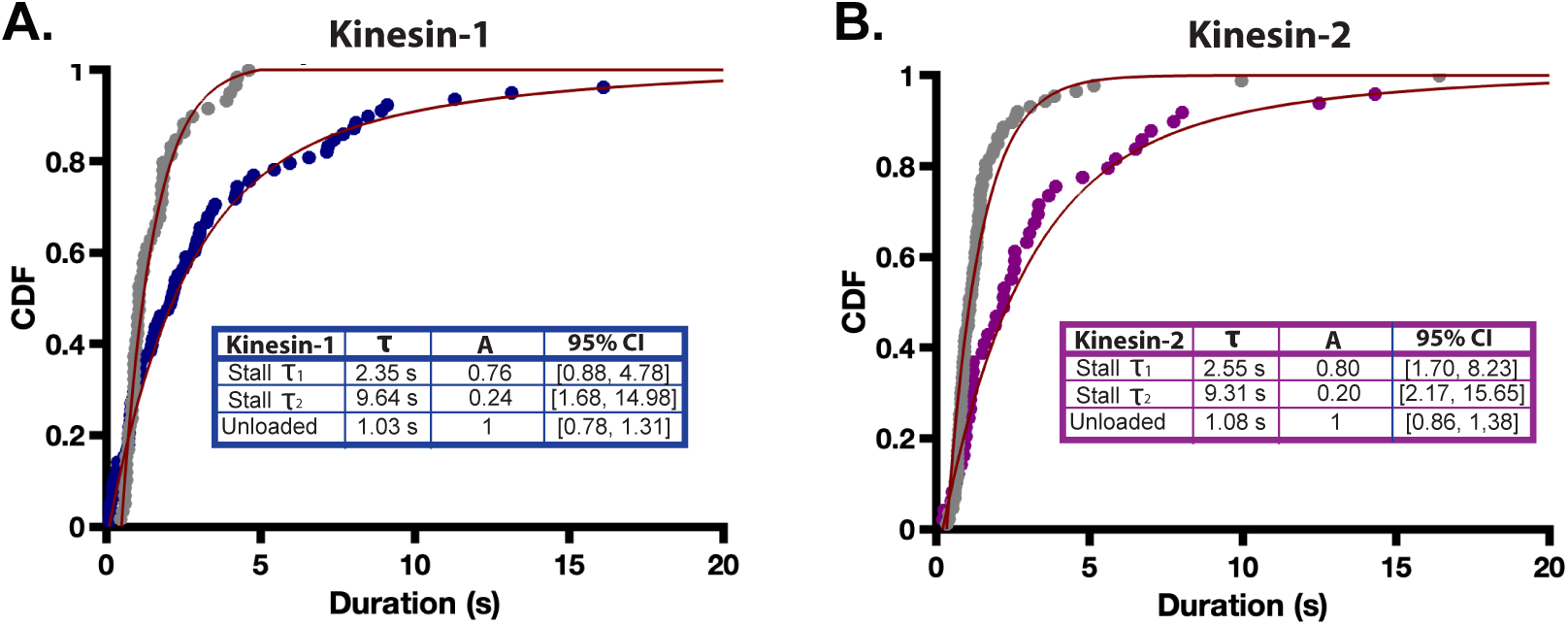
Bi-exponential fits of stall durations reveal a longer duration sub-population for kinesin-1 and kinesin-2. Tensiometer stall durations of (A) kinesin-1 and (B) kinesin-2 were fit with a biexponential function using a maximum likelihood estimator, MEMLET (https://michaelswoody.github.io/MEMLET/). Unloaded run durations are shown in gray for reference. Time constants (τ), relative amplitudes (A) and 95% confidence intervals for time constants are given in the accompanying tables. The rationale for why the motors would have two time constants is not clear, but it may suggest two alternative detachment pathways. Notably, both time constants are longer than the unloaded binding duration for both motors. Kinesin-3 stall durations were well fit by a single exponential function (see Fig. 2C).

**Figure 2 – figure supplement 2.**
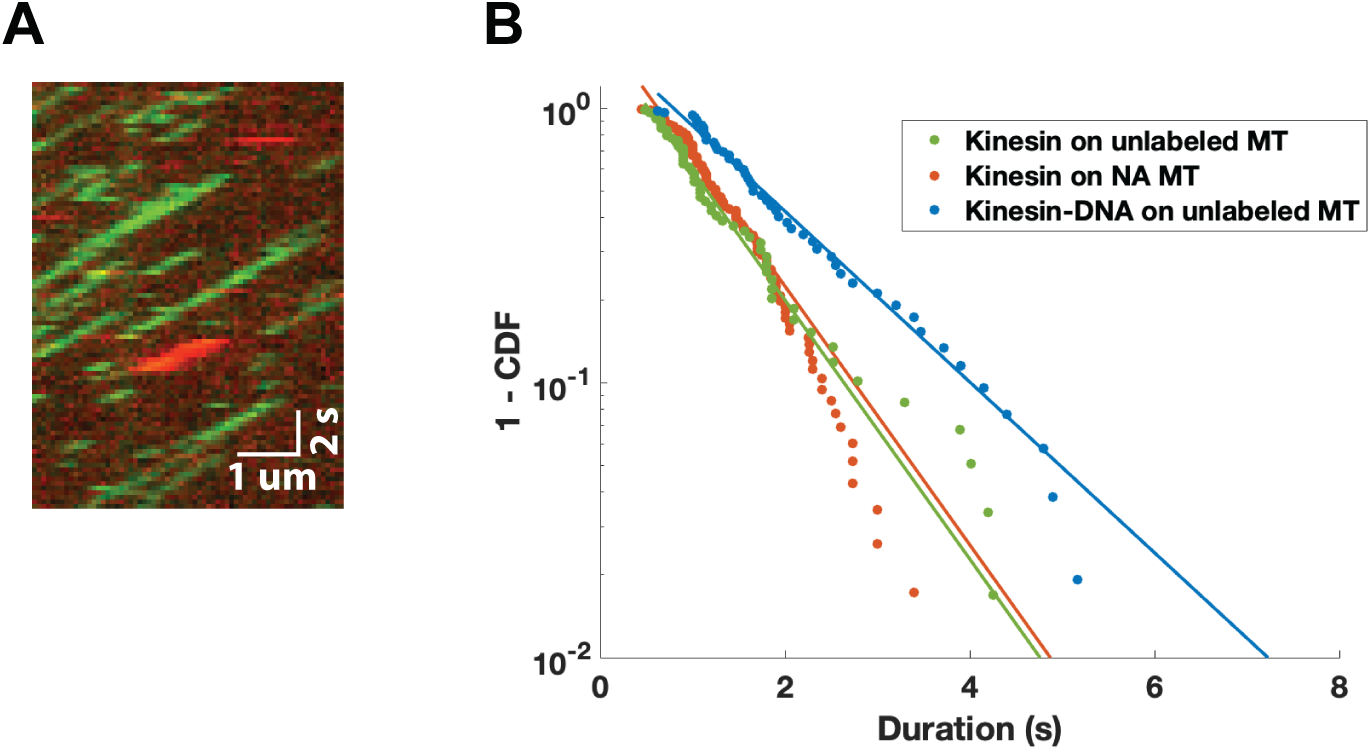
Kinesin-1 control experiments. (A) Kymograph of GFP-labeled kinesin-1 motors (green) transporting 3 kb Cy5-labeled dsDNA (red) along an unlabeled microtubule. Most motors do not have DNA attached, the one with DNA attached appears qualitatively similar to others. At top right, horizontal red line denotes free Cy5-labeled DNA diffusing near the microtubule for one frame. (B) Run durations for control experiments. Green circles: Unloaded GFP-labeled kinesin-1 on unlabeled microtubules; reproduced from Fig. 2A. From MEMLET fit, τ [95% CI] is 1.04 s [0.78, 1.31]. Red circles: Unloaded GFP-labeled kinesin-1 on biotinylated microtubules functionalized with neutravidin. Data were taken from excess non-DNA bound motors in DNA tensiometer experiments, thus matching the microtubules used for stall duration measurements. τ [95% CI] is 0.92 s [0.79, 1.04]. The similar run duration indicates that differences seen between unloaded run durations on unlabeled microtubules and durations of ramps and stalls in DNA tensiometer (on biotin-neutravidin functionalized microtubules) should not be due to effects of biotin-neutravidin. Blue circles: GFP-labeled kinesin-1 attached to 3 kb Cy5-labeled dsDNA moving on unlabeled microtubules (example shown in panel (A)). τ [95% CI] is 1.56 s [1.23, 1.90]. Thus, kinesin-1 transporting a 3 kb dsDNA has ∼50% longer run durations than kinesin alone. We interpret this difference to be due to the slower diffusion of the dsDNA, which enables undetected motor rebinding events that elongate the run length. Note that during ramps and stalls, any transient motor detachments should be detectable by a recoil of the stretched dsDNA, minimizing the effect of this slower diffusion of the dsDNA.

**Figure 4 – figure supplement 1.**
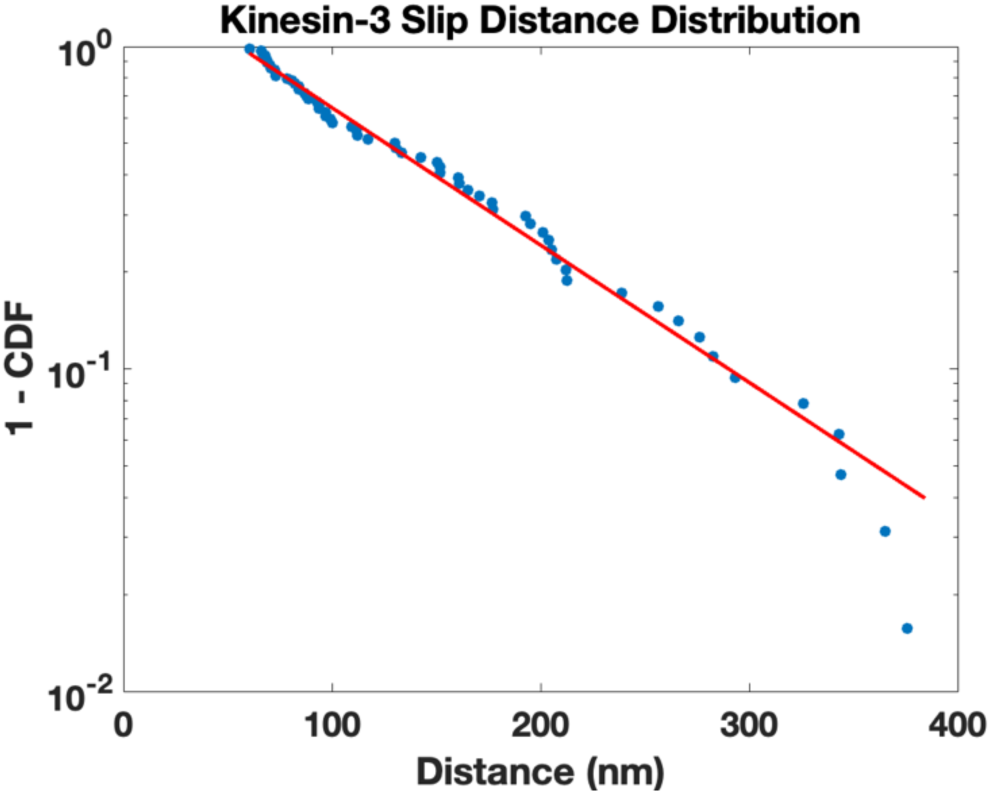
Correcting kinesin-3 stall duration for undetected slips. Because of the frequent slips during kinesin-3 stalls, we asked whether undetected slips below our 60 nm threshold might be leading to an overestimation of the kinesin-3 stall duration. To estimate the fraction of kinesin-3 slip events below the 60 nm threshold, we fit an exponential function to the distribution of slip events > 60 nm at stall:

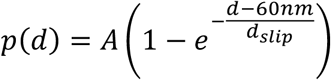 Because the data diverged at long distances, we only fit up to 400 nm and obtained d_slip_ = 102 nm (95% CI: 98-106 nm). The estimated fraction of slip events larger than 60 nm is: 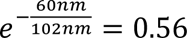, meaning that an estimated 44% of slip events were missed due to the 60 nm detection limit. Thus, the number of slip events should be adjusted upwards by 1/0.56 = 1.79. From the distribution of recovery times in Fig. 4F, 53% of plateau terminations were slip events (defined as the population with a 30 ms time constant) with the other 47% defined as detachment events. To correct for the total number of termination events, taking into account the theoretical missed slip events, we calculated a correction factor:

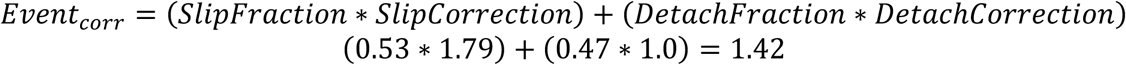 Thus, if undetected slips are taken into account, the number of stall termination events for kinesin-3 should be corrected upwards by a factor of 1.42, or equivalently the estimated stall duration of 1.89 s should be corrected to: 1.89 s/1.42 = **1.33 s.** Based on the d_slip_ fit uncertainty (98-106 nm), the 95% CI is 1.30-1.35 s. We note that this corrected kinesin-3 stall duration of 1.33 s is still well above the 0.75 s (95% CI: 0.64-0.87 s) for the ramp duration parameter in Fig 3 (Table S1). Thus, the kinesin-3 stall duration is longer than the kinesin-3 ramp duration, meaning that force slows detachment. There were only a handful of slips from stall for kinesin-1 and kinesin-2, precluding estimation of the fraction of missed slip events for these motors. However, the fraction is expected to be small and to not affect our conclusions.

**Figure 4 – figure supplement 2.**
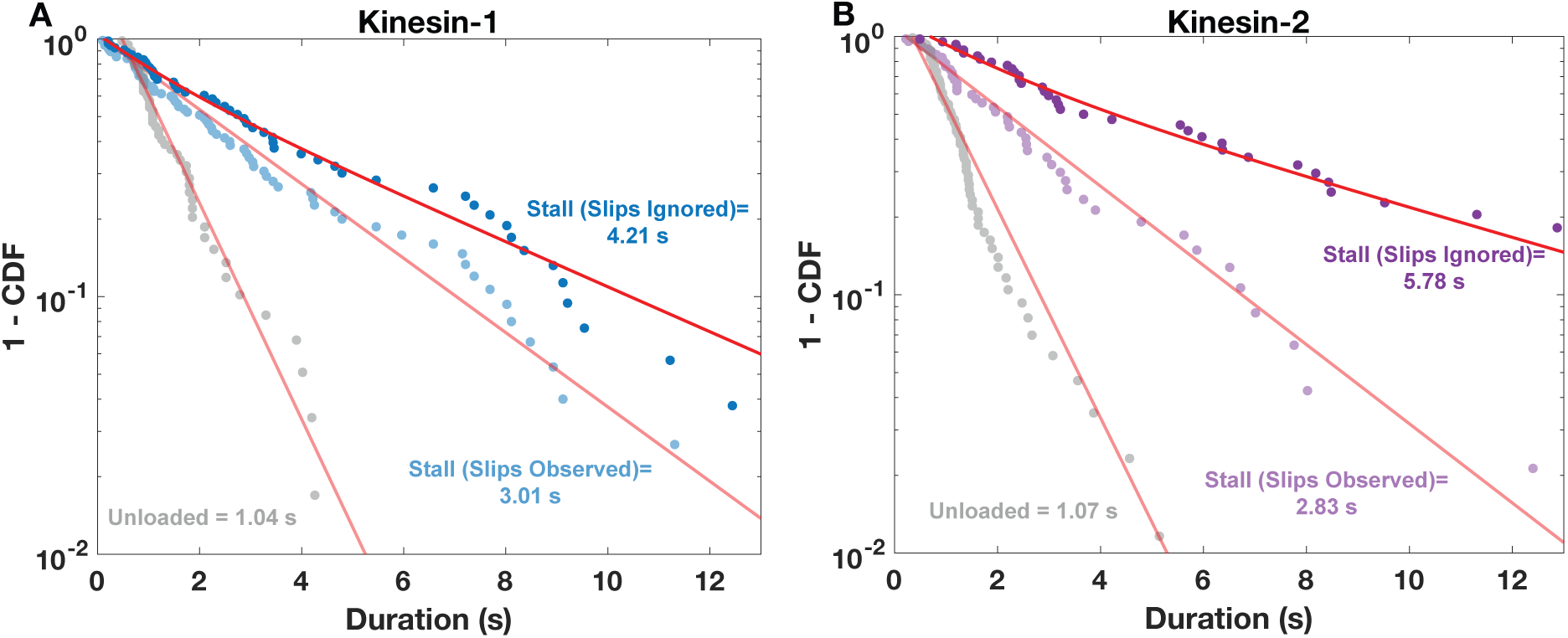
Stall durations are longer when slips are ignored. (A) Kinesin-1 stall durations, with unloaded run times in gray, stall durations terminated by slips in light blue, and stall durations terminated by falling to the baseline (ignoring slips) in dark blue. Unloaded and stall durations (from Figure 2) were fit with single exponential functions in MEMLET. Stall durations ignoring slips were fit with a bi-exponential by least squares τ_1_ = 1.35 s [95% CI: 0.81, 1.91 s], A_1_ = 0.27 s [0.07, 0.62 s], τ_2_ = 4.99 s [4.45, 5.93], A_2_ = 0.77 [0.46, 1.00]. Weighted average of the two time constants is displayed in plot for comparison to other time constants. (B) Kinesin-2 is plotted similarly in purple, fitting of stall duration ignoring slips resulted in τ_1_ = 1.91 s [95% CI: 0.92, 2.78 s], A_1_ = 0.32 s [0.11, 0.79 s], τ_2_ = 7.60 s [6.74, 9.05], A_2_ = 0.68 [0.41, 1.00].

**Figure 5 – figure supplement 1.**
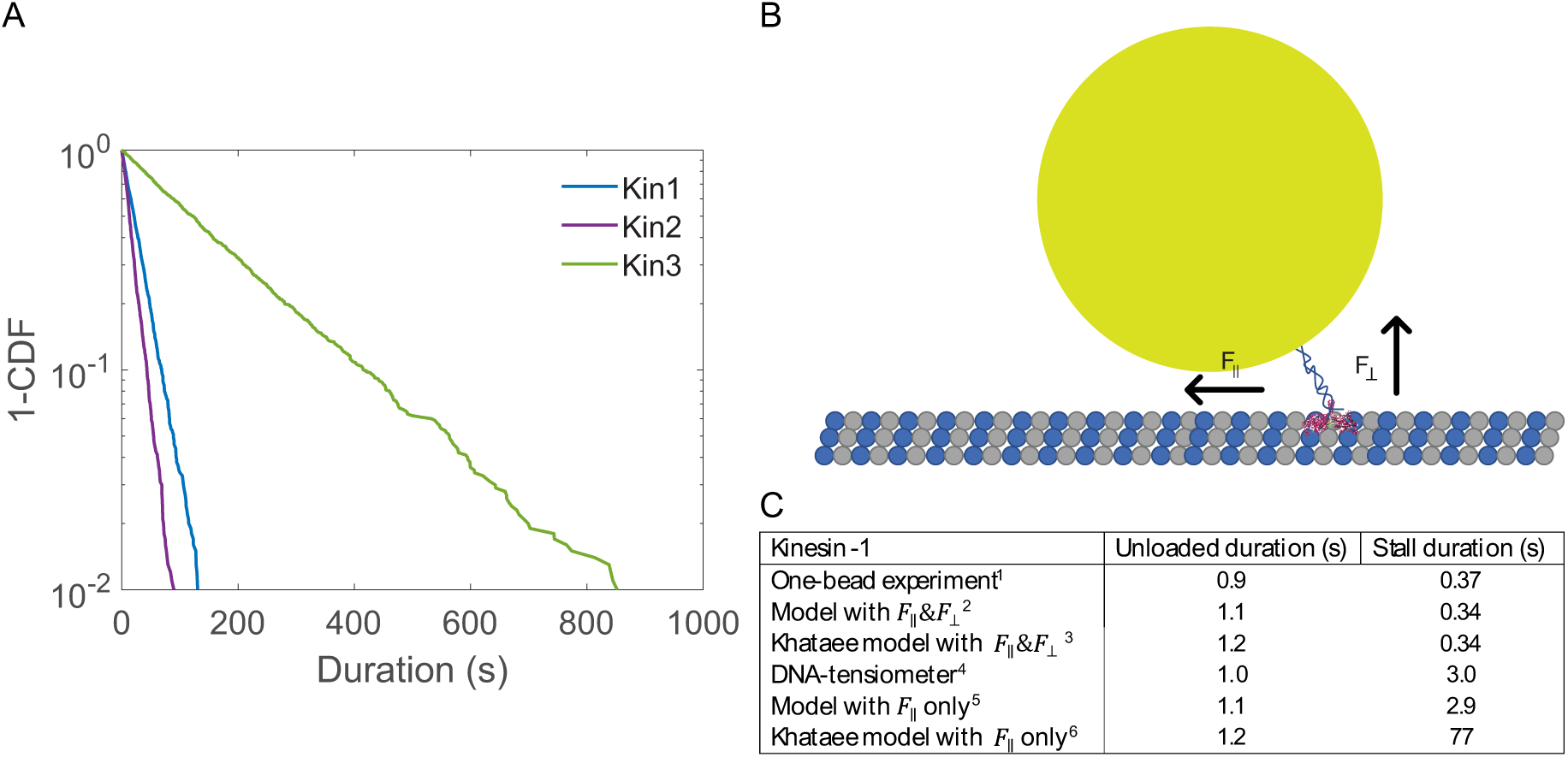
Model results incorporating both vertical and horizontal forces. (A) Model results incorporating load-dependent k_S-W_ and load-independent k_slip_. Very long stall durations are predicted that don’t agree with results, which motivated incorporating a load-dependence into k_slip_ (Figure 5). (B) Diagram of single-bead optical tweezer experiment using 440 nm diameter bead and 35 nm long motor, which results in a 60° motor angle. With this geometry, a 6 pN stall force parallel to the microtubule is associated with a 10 pN vertical force on the motor. Note that figure is not to scale. (C) Evaluation of effects of vertical and horizontal load from different models. ^1^One-bead experimental data at zero and 6 pN from Andreasson ^18,19^ ^2^Our model incorporating both horizontal and vertical forces into k_slip_ (using *δ*_‖_ = 1.61 nm and *δ*_┴_ = 1.58 nm) is able to match the single-bead stall duration. ^3^Khataee model ^19^ also accounts for single-bead unloaded and stall durations. ^4^Kinesin-1 DNA tensiometer data. ^5^When modeling our DNA tensiometer data for kinesin-1, our model using only F_‖_ is able to match the stall duration, whereas the Khataee model^6^ overestimates the stall duration.

## Notes

### Competing Interest Statement

The authors have declared no competing interest.

### Summary of Updates

Clarified text, reduced abstract to under 200 words, reorganized supplemental figures.

